# PTBP1 promotes cardiac hypertrophy and diastolic dysfunction by modulating alternative splicing

**DOI:** 10.1101/2020.06.30.171983

**Authors:** Carlos Martí-Gómez, Javier Larrasa-Alonso, Marina López-Olañeta, María Villalba-Orero, Pablo García-Pavía, Fátima Sánchez-Cabo, Enrique Lara-Pezzi

## Abstract

Alternative splicing (AS) plays a major role in the generation of transcript diversity. In the heart, roles have been described for some AS variants and individual regulatory RNA binding proteins (RBPs); however, the global impact and regulation of AS patterns in cardiac pathophysiology is poorly understood. Here, we studied the AS profiles in heart disease, their relationship with heart development and the regulatory mechanisms controlling AS dynamics in the mouse heart using a total of 136 RNA-seq samples. We found that AS and gene expression changes affect different genes, which are also involved in distinct biological functions. Developmental AS changes were more abundant and had stronger predicted impact on the encoded protein than those taking place during heart disease. However, AS changes in heart disease significantly modified protein interaction patterns and rewire the protein-protein interaction network. Using a database of experimentally determined binding sites of a large collection of RNA binding proteins, we studied the regulatory proteins associated to AS changes in each condition. Computational modelling revealed that developmental transitions were mainly driven by the up-regulation of MBNL1, whereas disease associated AS changes were driven by a more complex regulatory network, characterized by the interaction of different RNA binding proteins, with PTBP1 as the largest individual modulator. In adult mice, PTBP1 over-expression was sufficient to induce cardiac hypertrophy and diastolic dysfunction and significantly alter the AS profile. Overall, our study provides new insights into the functional impact of AS patterns in cardiac physiology and how computationally driven hypotheses can help to improve our understanding of RNA regulation and its contribution to heart disease.

## Introduction

Cardiovascular diseases are the leading cause of mortality and morbidity worldwide. In the USA, the prevalence of coronary heart disease is 6.3% among adults aged above 19 years and accounts for more than half of cardiovascular events in individuals < 75 years old (7). Aortic stenosis has a prevalence of 0.4% in the entire US population, rising to 2.8% among the elderly. Despite improvements in our knowledge about gene expression (GE) patterns, understanding of the molecular mechanisms underlying the development of heart disease remains incomplete (41). Specifically, there is poor understanding of the role and regulation of different transcript isoforms generated by Alternative splicing (AS).

AS enables the generation of different transcripts from a single gene (3). Comparative studies suggest that AS might be particularly important for the function of brain, heart, and skeletal muscle (54). In the heart, regulation of AS during postnatal development is important for proper fiber maturation, including the regulation of passive stiffness by differential inclusion of titin exons (39). Many disease-causing mutations affect AS, further supporting its physiological relevance (41). For example, myotonic distrophy (MD) is caused by expansion of CTG repeats in the DM protein kinase (DMPK) 3’ UTR. These sequences sequester MBNL1 and preclude its binding to other targets for AS regulation (36).

The study of the AS landscape and its regulatory mechanisms in the mouse heart has mainly focused on developmental stages, in particular in the postnatal transition (1, 26, 35, 73). In addition, although AS of each exon can be controlled simultaneously by multiple RNA binding protein (RBP), previous analyses have focused on single regulatory proteins (mainly the Mbnl and Celf families) acting individually (2). Little is known about AS regulation at earlier stages of development or in heart disease. In addition, although heart development and heart disease are assumed to be highly related, the mechanisms underlying reexpression of the neonatal AS profile in heart disease remain mostly unknown (2). Most transcriptomic studies have been performed with a limited number of biological replicates (26), with little to no representation of interindividual or interlaboratory variability and therefore limiting the interpretation of the results. Here, we integrated data from 21 published RNA-Seq experiments, with a total of 144 samples covering embryonic, neonatal, and adult heart, and 2 widely used mouse models of heart disease: pressure overload cardiac hypertrophy, induced by trans-aortic constriction (TAC), and myocardial infarction (MI), induced by permanent ligation of the left anterior descending coronary artery (LAD). Appropriate statistical analysis of this large dataset allowed us to identify changes between conditions and to account for different sources of variability. We then used these well characterized phenotypes to study the functional relevance of AS across development and disease and to determine the underlying regulatory mechanisms.

## Results

### Characterization of AS patterns in the developing and diseased heart

To characterize the AS changes taking place during heart development and disease, we collected a large dataset of heart samples from mouse models at different developmental stages and disease conditions (Table S1). We grouped the collected samples according to 5 major phenotypes, including the major previously characterized developmental stages: embryo (E10.5-E17), post-natal (P0 to P7), and adult (from P10) (1, 26, 35) and two common models of heart disease: TAC and MI. Using this categorization of samples, we first identified a set of over 20,000 AS events with at least one read supporting skipping or inclusion in 20% of the samples. We then used a series of Generalized Linear Mixed Models with binomial likelihood to identifyAS changes occurring in 4 specific transitions: embryonic development (ED), by comparing neonatal with embryonic samples; post-natal development (PD), by comparing uninjured adult samples with neonatal samples; TAC, by comparing samples from hypertrophic and uninjured adult hearts; and MI, by comparing samples from infarcted and uninjured adult hearts. We incorporated sample and experiment as random variables to take into account biological variability in inclusion rates and batch effects, respectively.

AS changes were more abundant in the developmental transitions than in the disease models, suggesting more prominent roles of AS during embryonic and postnatal development (Figure 1A, Table S2). Given that the main AS changes in all comparisons were cassette exon events, we focused on this type of event in downstream analyses. Interestingly, whereas exon inclusion and skipping were observed in similar amounts during both embryonic and post-natal development, heart disease was mainly characterized by increased exon skipping (Figure 1).

**Fig. 1.**
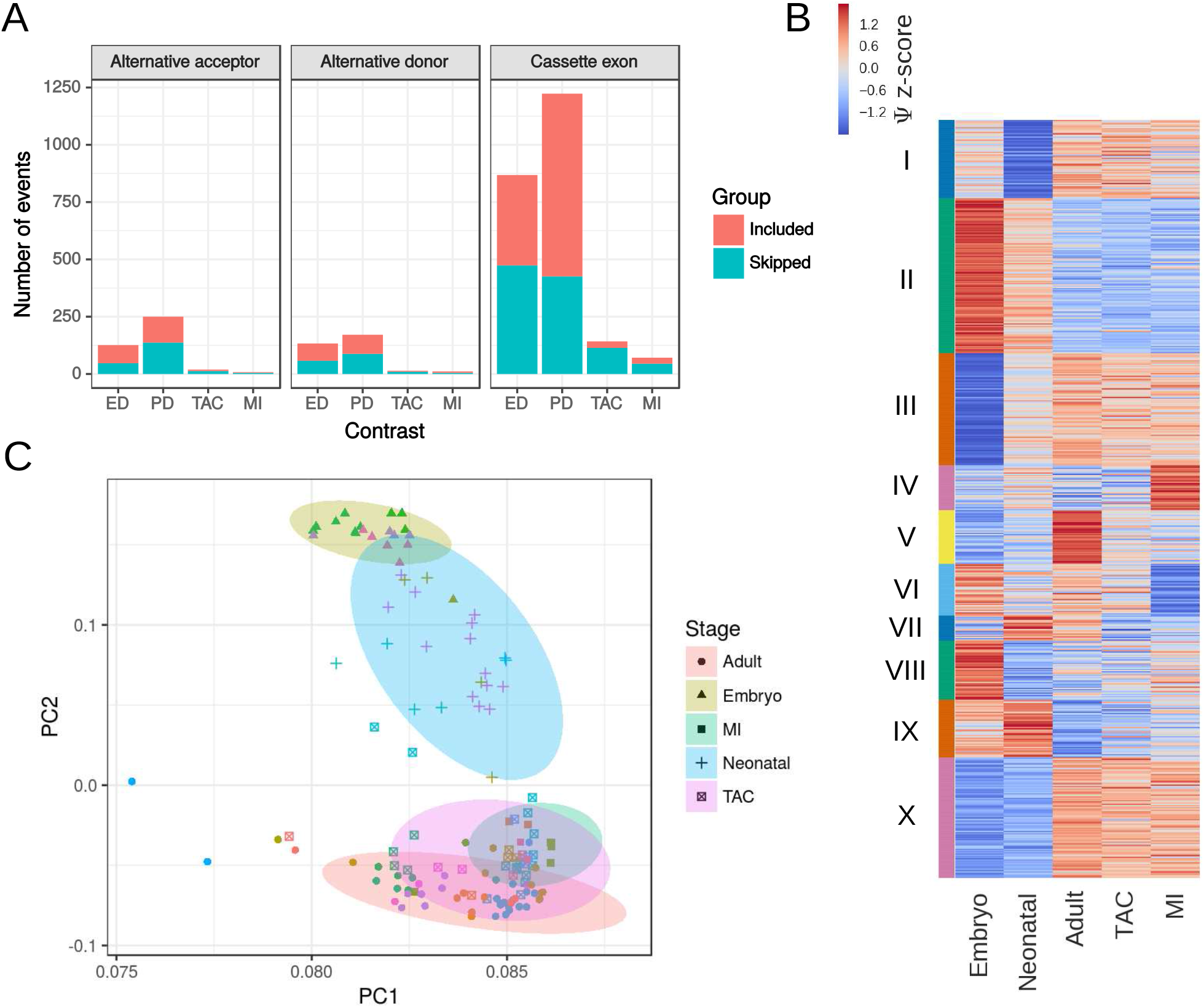
Alternative splicing landscape in heart development and disease. **A** Number of events showing significant differences in each comparison according to the event type: alternative acceptor, alternative donor, and exon skipping. **B** Heatmap representing z-scores calculated from estimated Ψ for each condition. Per condition Ψ was estimated using a linear mixed model with binomial likelihood function. Clusters were calculated using k-means on the normalized Ψ profiles. **C** Principal Component Analysis of all analyzed samples using exon cassette events without missing data in any of the samples. Different symbols represent different conditions and colors represent different datasets or experiments. Ellipses were drawn according to each condition automatically using geom_ellipse function from ggplot2 library (75).

In agreement with the observations in Figure 1A, K-means clustering of the standardized Ψ profile (Figure 1B, Table S3) revealed that the largest clusters (II,III,IX,X) were those specific of developmentally regulated exons, with smaller clusters identified with specific changes in TAC and MI (clusters IV and VI). Interestingly, clusters V, VII and VIII show similar pattern in MI and in embryonic samples, suggesting partial re-expression or re-repression of the neonatal AS pattern after cardiac injury. Furthermore, Principal Component Analysis (PCA) showed a small displacement of TAC and MI samples toward neonatal samples (Figure 1C) reinforcing this idea. Embryonic and neonatal samples were clearly separated from the adult samples, mostly on PC2 but to some extent also on PC1, which harbors 93% of the global variance. We did not observe a particular association of batches over the main PCs, suggesting that even if there is a lot of variability within groups in PC1, there are AS signatures that differentiate the different groups under study. This was, nonetheless, not specific to AS, as PCA of expression data showed a similar pattern (Figure S1, Table S4).

### AS changes modulate low impact exons and are independent of GE changes

We next investigated the potential impact of AS changes on heart physiology. Two of the main potential roles of AS are the production of different protein isoforms from the same gene and the regulation of gene expression by regulated unproductive splicing and translation (RUST). Exons that generate alternative protein isoforms usually preserve the ORF to avoid early termination of protein translation. Significantly changed exons had more often lengths divisible by 3, and thereby were more likely to preserve the reading frame upon either inclusion or skipping (Figure 2A; Fisher test p-value <0.01 for all comparisons except MI Included and TAC Included). Moreover, exons preferentially included in developmental transitions and those preferentially skipped in TAC or MI were shorter in average than those that showed no significant changes (Figure 2B, MannWhitney U test p-value < 0.05). These exons, which were the most abundant (Figure 1A), showed an under-representation of PFAM domains. In contrast, a higher proportion of exons skipped in the ED or PD comparisons or included in MI encoded PFAM domains (Figure 2C, Fisher test p-value < 0.05). These results suggest that quantitative AS changes in the heart occur mainly in coding regions, but are predicted to have only minor consequences for protein function.

**Fig. 2.**
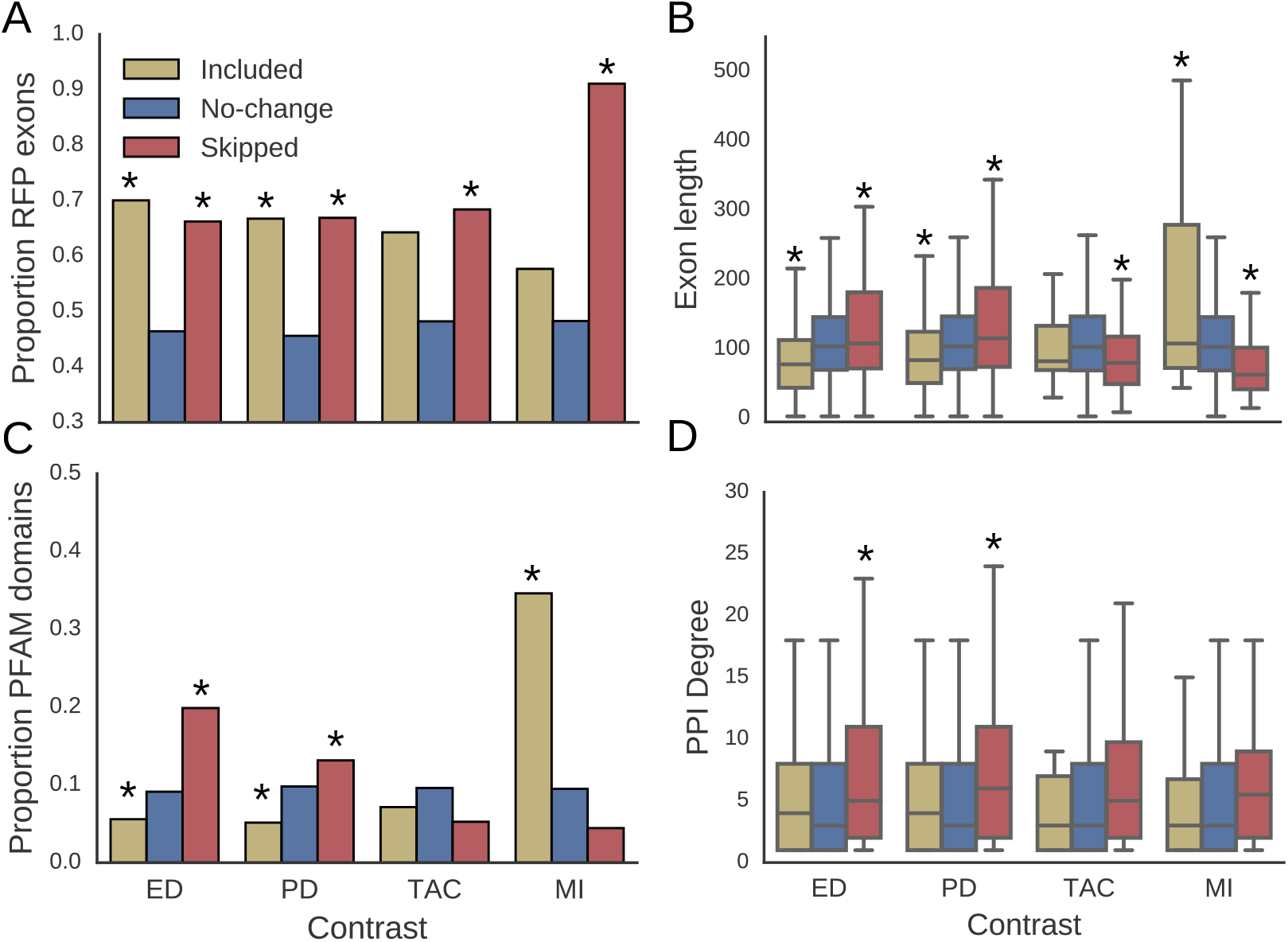
Properties of the 8892 alternatively spliced exons under study in the heart. **A-D**, Values of different properties in each comparison according to whether inclusion levels where increased (Included), decreased (Skipped), or not significantly changed (No-change). A total of 394, 797, 28 and 26 exons were included in ED, PD, TAC and MI, respectively; whereas 474, 426, 114, 45 exons were skipped. **A** Proportion of exons with length that is multiple of 3 (reading frame preservation (RFP)) and therefore have no impact on the open reading frame. **B**, Exon length distribution. **C**, Proportion of exons overlapping with an annotated PFAM domain. **D**, Number of connexions or degree in the Intact protein-protein interaction (PPI) network. Differences in proportions between Included and Skipped groups compared with No-change were assessed using Fisher exact tests; differences between Included and Skipped groups compared with No-change in exon length and PPI degree, were tested using Mann-Whitney U tests (* represents p-value<0.05).

We next investigated the overlap between changes in AS and changes in GE. The proportion of differentially expressed genes was similar in genes undergoing differential AS and in those showing no AS changes, suggesting no association between AS and GE changes (Figure 3A). Even if GE and AS regulate different genes, these affected genes might regulate the same biological processes. Thus, we performed Gene Ontology (GO) enrichment analysis in each comparison and then calculated the pairwise semantic similarity of the 10 most significantly enriched GO terms among all groups followed by hierarchical clustering based on the similarity profile (Figure 3B). This analysis clustered enriched processes for differentially spliced and differentially expressed genes separately, regardless of the biological context (development or disease), indicating that AS and GE changes affect distinct biological processes in the heart. Interestingly, processes regulated by AS clustered separately for disease and development, whereas processes associated with GE changes clustered together (upregulated in development and downregulated in disease clustered separately from downregulated in development and upregulated in disease). This suggests a stronger functional reexpression of embryonic GE patterns than AS patterns in heart disease. Whereas changes in GE were mainly related to cell division, the respiratory chain, and extracellular matrix deposition, AS changes were more associated with cytoskeletal organization (Figure 3C).

**Fig. 3.**
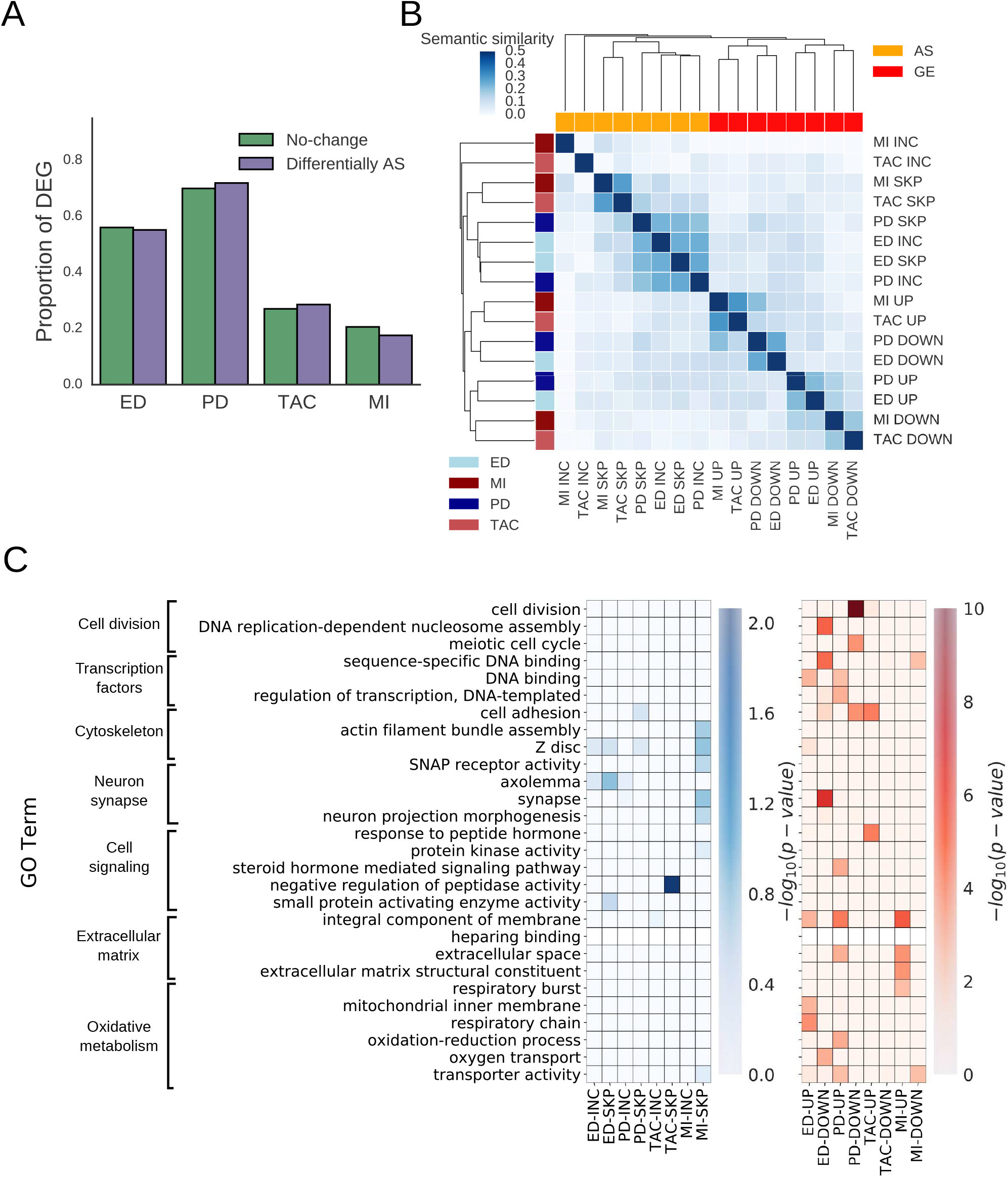
Functional impact of AS and GE changes and their overlap. **A**, Proportion of genes showing GE changes that also show changes in AS in each comparison. Differences in proportions were tested using a Fisher exact test using a total of 4697 alternative genes, out of which 619, 897, 127 and 58 genes showed significant AS changes in ED, PD, TAC and MI, respectively (p-value > 0.1 for all contrasts) **B**. Pairwise semantic similarity among the most representative GO terms in each group of genes. Row colors represent the different comparisons studied and column colors represent AS or GE. Semantic similarity profiles were then clustered using hierarchical clustering using euclidean distance and Ward method for grouping. **C**, Heatmaps representing the –log10(p-value) of the functional enrichment analysis for top enriched categories across all groups for AS (left) and GE (right). Independent categories were selected with an L-1 regularized logistic regression first, followed by a regular logistic regression to account for all gene categories simultaneously. P-values refer to the significance of the coefficient representing each gene category in the regression model (see Methods for details)

### Modulation of protein-protein interaction networks by AS in heart disease

Alternative splicing protein isoforms have been previously shown to have different interaction partners (77). To investigate whether AS changes in the heart regulate protein protein interactions we first compared the connectivity of proteins encoded by genes undergoing differential AS using the Intact Protein-Protein Interaction (PPI) network (58). For all comparisons, genes with skipped exons showed more connections to other proteins than genes without significant AS changes (Figure 2D; Mann-Whitney U test p-value < 0.05 for PD and ED, p-value < 0.15 for MI and TAC). To check whether this was a general property of transcriptional changes, we compared the PPI degree distribution of proteins depending on their gene expression changes. In contrast to AS, DEGs did not show a higher number of connections in the PPI network (Fig. S2), suggesting that this feature is specific to AS. However, this does not necessarily mean that these particular AS changes are modulating the interaction capabilities of these proteins. To determine whether PPIs are actually regulated by AS changes, we used information of Domain-domain interaction (DDI) information (24), and assumed that exons located in a domain that mediates an interaction between two proteins are actually required for such interaction to take place. Thus, we can identify interactions that are increased or decreased depending on whether the inclusion of the exon spanning the domain increases or decreases, respectively. We found that exons included during TAC or MI affected more domain mediated PPIs than unchanged exons (OR=3.50 and OR=2.42, Fisher tests p=0.06 and p=0.21, respectively). Skipped exons in disease, if anything, avoided changing interactions (OR=0.51 an OR=0.65, p-values > 0.2) (Figure 4A). Overall, these results suggest that AS changes can increase the number of interactions by increasing exon inclusion, but avoid reducing them through exon skipping.

**Fig. 4.**
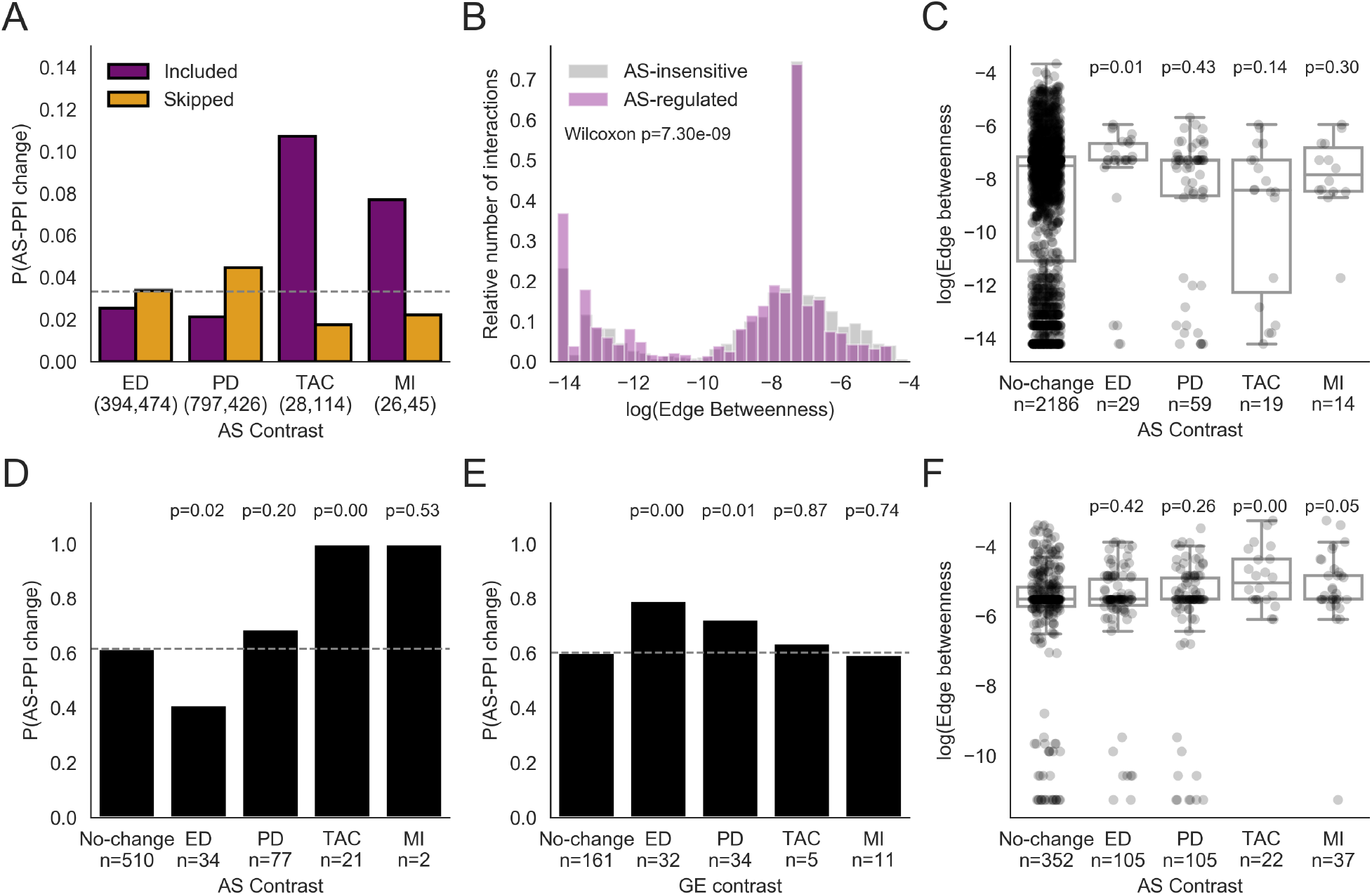
Impact of AS changes in the protein-protein interaction networks. **A**. Estimated proportion of exons that map to a domain mediating protein-protein interaction according to whether they are included, skipped or unchanged in the comparisons under study. Numbers in brackets represent the number of exons in the groups Included and Skipped for each contrast, out of 8893 total exons under study. **B**. Distribution of the log(Edge betweenness) for interactions that were found to be potentially regulated by AS in the heart compared to the remaining interactions. An interaction was considered to be potentially modulated by AS when an exon considered to be alternative in the heart encoded a domain mediating this interaction. **C**. Network edge betweenness for interactions depending on whether they were modified by AS in the conditions under study using domain mediated interactions (24). **D,E**. Estimated proportion of interactions that differed between AS isoforms (76) in genes with significant differences in AS (D) or GE (E) in each contrast. **F**. Network edge betweenness for interactions depending on whether they were modified by AS in the conditions under study using data in (76). Fisher tests were used to assess differences in proportions and Mann-Whitney U tests to analyze differences in edge betweenness

The impact of increasing or decreasing the amount of interacting proteins may however depend on their position in the PPI network. In other words, reduction or even complete ablation of the binding affinity between two proteins by differential exon inclusion may only affect slightly the overall function of a protein complex, as cooperative binding of the remaining elements may compensate this lack of binding between some of their elements. On the other hand, if two protein complexes interact only through an interaction between a pair of proteins, the modulation of this interaction is expected to have greater functional consequences. Thus, we analysed how AS changes affect the structure of the PPI network beyond isolated interactions. To do so, we built an undirected graph using pairwise interactions for genes expressed in at least one condition and calculated the betweenness for each edge. DDI interactions potentially modulated by AS showed, in general, lower betweenness, compared to AS-insensitive interactions (Figure 4B, MannWhitney U test p-value< 10^−6^). These results suggest that AS-modulated interactions tend to be located within closely interacting modules rather than connecting different protein complexes. When comparing across groups of modulated exons, we found that exons modulated during ED have significanly higher betweenness in the PPI network than the unchanged exons (Mann-Whitney U test p=0.01, Figure 4C), suggesting a stronger rewiring of the interaction networks during early heart development than in any other condition. No significant difference was found in the betweenness for exons modulated during heart disease (Mann-Whitney U test p>0.1, Figure 4C).

Since proteins do not only interact through protein domains, we next studied AS-mediated PPI changes in experimentally built networks that are not limited to DDIs (76). We found that 100% of exons changing in disease AS changes are located in genes with known AS-dependent interactions (Figure 4D), significantly greater than the approximately 60% observed for unchanged exons (Fisher test p<0.0001 and p=0.53 for TAC and MI, respectively). These findings are specific to AS since GE changes showed the opposite trend: only developmentally regulated genes are associated to AS-modulated interactions (Figure 4E). To investigate the global impact of AS on the PPI network, we built an interaction network using only experimentally tested interactions in this dataset and calculated the edge betweenness, as before. Interactions affected by AS changes in TAC and MI showed higher betweenness than unchanged exons (Mann-Whitney U test p<0.01 and p=0.05, respectively, 4F), suggesting a rewiring of the PPI network by AS in heart disease. Despite the low statistical power due to the small size of groups overlapping with available PPI data in each dataset, our results suggest that AS changes significantly alter PPIs networks in heart disease.

### RBPs associated to AS changes during development and disease

To identify the potential regulators of AS in the heart, we looked for over-represented binding sites of different RBPs across different potential regulatory regions. Binding sites were collected by integrating a series of databases of CLiPseq experiments (see Methods). We first filtered those RBPs that were found to be significantly enriched (p<0.01, Fisher test) in at least one group of significantly changed exons. We then used the reduced set of enriched RBPs binding to different regulatory regions as substrate for regression analysis using a GLM with binomial likelihood to take into account co-linearities across binding profiles of different RBPs. This analysis was then applied to sets of exons that were found to change in any comparison (Figure 5A). Our results show that MBNL1 is strongly enriched in the upstream intron of exons that are skipped and in the downstream intron of exons that are included during both PD and ED. We also found that MBNL1 binding sites in exons showing changes tend to be more conserved across evolution at the sequence level, suggestive of functional importance (Figure 5B). Additionally, MBNL1 expression increases during development and remains unchanged in both TAC and MI (Figure 5C). To test whether different RBPs may regulate different biological functions, we looked for enrichment of GO terms in exons bound by each RBP compared to all those that changed in any of the comparisons under study (Figure 5D, one-tail Fisher test). We found that MBNL1 tends to bind to genes related with actin cytoskeleton dynamics and cell junctions, whereas other RBPs tend to bind more to exons of RNA binding proteins or proteins located in the nucleus. Whereas other RBPs may contribute to the regulation of AS changes during development, such as QK, RBFOX1 or PTBP1/2, our results suggest that MBNL1 is the main regulatory element.

**Fig. 5.**
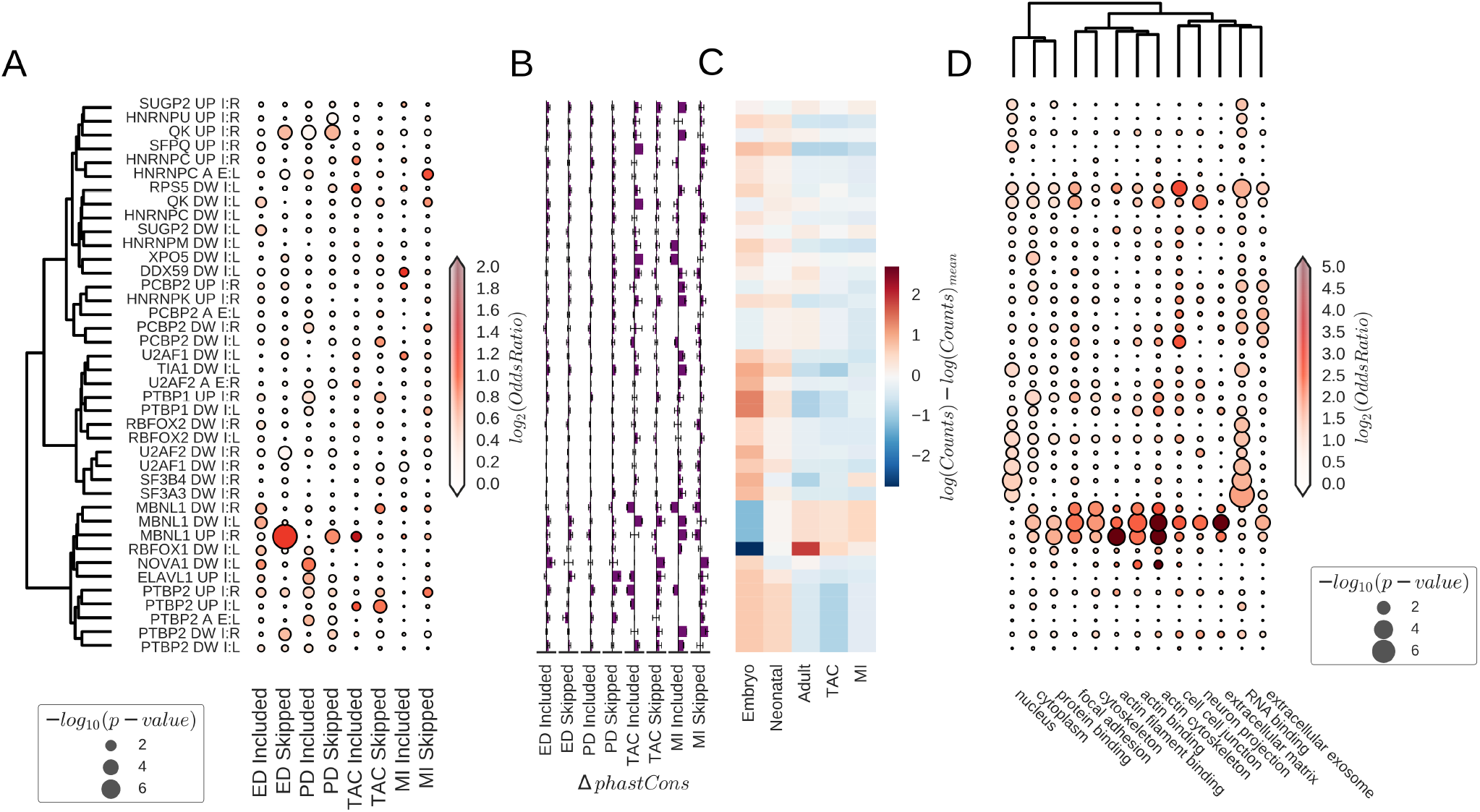
Direct regulation of alternative splicing changes. **A** Dotplot representing multivariate enrichment analysis for CLiP-seq binding sites in regulatory regions for RBPs that showed significant enrichment by univariate analysis for each group of exons. Regions are defined by combinations of the following terms: (I: Intron, A: Alternative exon, DW: downstream, UP: upstream, L: left, R: right). The dendrogram was calculated using the Ward method with distances between binding sites profiles to the exons included in the analysis. **B** Mean difference in phastCons scores in binding sites of exons showing changes compared to those that remained unchanged. Error bars represent the standard error of this difference. X-axis range from −0.5 to 0.5 in all cases. **C** Centered expression levels per condition under study for RBPs showing significant univariate enrichment from panel. **D** Dotplot representing the functional enrichment of genes with binding sites for each RBP and region and showing significant changes in at least one comparison. Genes with significant AS changes in at least one comparison were used as background for enrichment (one-sided Fisher test). The dendrogram represents distances between GO terms based on the proportion of shared genes using the Ward method.

Although MBNL1 was found to be the main regulator of AS during ED and PD, we found only a mild enrichment of MBNL1 binding sites in changing exons in both models of heart disease: TAC and MI. In addition, MBNL expression remained unchanged in disease, suggesting that MBNL1 does not modulate AS changes in disease (Figure 5C). In contrast, not only PTBP1 expression decreased during development, but also increased upon MI or TAC (Figure 5C). PTBP1 and PTBP2 binding sites were enriched in the upstream intron of skipped exons in TAC and MI, and included in ED and PD (Figure 5A). All this suggests that the binding of PTBP1 to the upstream intron of alternative exon inhibits exon inclusion and that PTBP1/2 might mediate the partial re-expression of neonatal AS patters in heart disease.

### Reduced coordination of RBP expression changes is associated with complex regulatory mechanisms of AS in heart disease

Independent RBP enrichment analysis identified MBNL1 and PTBP1/2 as the main regulatory elements of AS during heart development and disease, respectively. However, the effect of one RBP may depend on the binding of another RBP, cooperative or competitive binding, giving raise to more complex regulatory patterns. To investigate the relative contribution of cooperative or competitive binding, we expanded our logistic regression model to include all pairwise combinations of RBP binding sites. As we expect most of these interactions to have no effect on the inclusion patterns, we added a Lasso penalization to promote sparsity of explanatory variables. We first optimized the regularizing constant using 10-fold cross-validation in terms of the AUROC, as a measure of predictive power (Figure 6A). Larger regularizing constants are required for TAC and MI models than for ED and PD, showing that they require a larger set of non-zero regulatory activities to explain the observed changes. As expected, interactions were less likely to contribute to predict inclusion or skipping of a particular exon (Figure 6B), even if the effect sizes of single and pairs of RBPs were comparable (Figure 6C). However, since the number of possible RBPs pairs is higher than the number of RBPs, the cumulative contribution of pairwise interactions to modulate inclusion rates was comparable in ED and PD to that of single RBPs, and was even higher in MI and TAC (Figure 6D). This suggests that the underlying regulatory patterns of AS in heart disease are more complex than those modulating AS during development. Figure 6E and F show the specific non-zero interactions among RBPs for included and skipped exons in ED and MI, respectively (PD and TAC patterns show the same trend at Figure S3). Interestingly, we found that for MI and TAC (Figure 6F and S3B, barplot on the right), PTBP1 binding to the upstream intron showed the greatest contribution individually among all considered RBPs, suggesting again a prominent role of PTBP1 within the complex regulatory network underlying AS changes in disease.

**Fig. 6.**
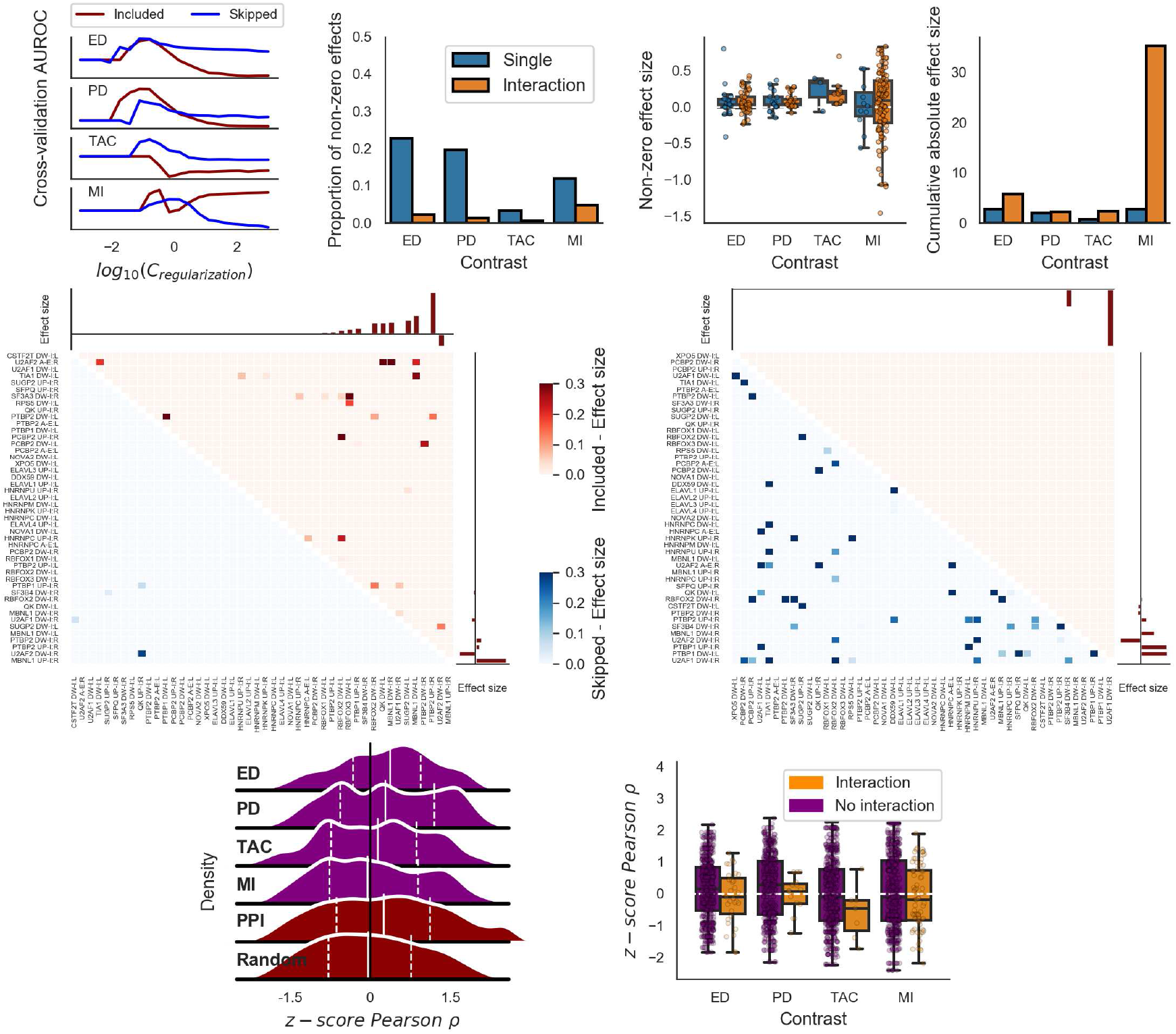
Analysis of AS regulatory complexity using regularized logistic regression with all pairwise combinations of RBPs binding sites. **A** area under the Receiver Operating Characteristic curve (AUROC) in 10-fold cross-validation analyses along a range of regularizing constants (*C_regularization_*; the lower, the stronger the regularization), used to select the value with strongest predictive power of a particular group of exons. **B, C, D** Proportion of non-zero estimations (B), coefficient estimates (C), and cumulative absolute values of coefficient estimates (D), for coefficients corresponding to a single RBP and to an interaction between a pair of RBPs for each comparison under study. **E, F** Heatmap representing the estimate of the coefficients for each combination of RBPs for exons that are either included (Red) or skipped (Blue) for ED (E) and MI (F). Barplots represent the estimation of the coefficient for single RBPs. **G, H** Distribution of normalized correlation coefficients between expression levels of RBPs included in the regression model for each comparison (ED, PD, TAC, MI) and for pairs of interacting proteins and randomly selected pairs of genes as a whole (G) and separating pairs that showed non-zero coefficient (H).

We hypothesized that this increased complexity arises from the lack of coordination in RBPs expression changes. If the expression of two RBPs increased or decreased while maintaining their stoichiometry, they would act mostly in complexes and only their common targets would change. However, if there was an imbalance in their expression levels, both common and individual targets of these RBPs would change. Therefore, if lack of coordination underlay this increased regulatory complexity, we would expect a lower correlation among RBPs in TAC and MI than in ED and PD. We found comparable normalized correlation coefficients among candidate RBPs in ED and PD to that of pairs of interacting proteins from the Intact PPI database (Figure 6G). The correlation was however lower in MI and TAC (linear regression model, p-value=0.016 and p-value=0.062, respectively). Furthermore, within each comparison, pairs of RBPs showing non-zero regulatory interactions at the binding site level showed lower expression correlation (Figure 6H, linear regression model, p-value=0.026). These observations, altogether, are consistent with a model with an altered stoichiometry of regulatory proteins that results in more complex regulatory patterns in disease conditions.

### PTBP1 over-expression induces cardiac hypertrophy and diastolic dysfunction

To further investigate the potential contribution of PTBP1 to AS changes, we used published RNA-Seq data from a PTBP1 and PTBP2 loss-of-function *in vitro* experiment in neural progenitor cells (13, 25, 42, 48) (Figure S4). AS changes in the PTBP1/2 double KD correlated with those in the cardiac PD, TAC, and MI contrasts. The stronger correlation with the double KD (r=-0.15) than with the individual PTBP1 KD (r=-0.087, Likelihood ration test p-value< 10^−16^) suggests that both PTBP1 and PTBP2 actively regulate AS in the heart in all the conditions studied, in agreement with the bindingsite–enrichment analyses (Figure 5). In contrast, when analyzing MBNL1 deficient hearts (25), we found a correlation only with developmental AS changes, further supporting that MBNL1 does not mediate AS changes in disease (Figure S4). After confirming that PTBP1 was upregulated in independent TAC and MI experiments by qPCR (Figure 7A,B), we aimed to investigate whether PTBP1 upregulation alone is sufficient to induce pathological changes in the heart. We over-expressed PTBP1 specifically in the heart using an Adeno-Associated virus type 9 (AAV9) as vector carrying PTBP1 cDNA under the control of a cardiac-specific promoter (TnIc). We injected AAV9-PTBP1 into 10-12 weeks old wild-type (WT) mice in two independent experiments, using luciferase-expressing AAV9 as a control, and analyzed the mice 28 days later. We found that PTBP1 over-expression in mice injected with AAV9-PTBP1 was similar to that observed in TAC experiments (1.2 fold, Figure 7C, linear regression model p-value=0.008). We evaluated cardiac function *in vivo* using transthoracic echocardiography 28 days post-injection. Mice over-expressing PTBP1 showed an increased normalized cardiac mass (Figure 7D, linear regression model p-value=0.001), particularly in the left ventricular posterior wall. Although no significant change in left ventricle ejection fraction was observed (Figure S5, linear regression model p-value=0.59), suggesting normal systolic function, we found a reduction in the E/A ratio (linear regression model p-value=0.017). This lower E/A indicates a de-compensation on the relative contribution of passive and active left ventricle filling or, in more general terms, diastolic dysfunction (Figure 7E). We also observed a significant increase in the expression of cardiac dysfunction markers MYH7 and BNP (p-value<0.01) following PTBP1 over-expression. We found no significant changes in the expression of fibrosis markers by qRT-PCR (LOX, COL1A1, COL3A1, linear regression model p-value>0.45) (Figure 7F) or in collagen quantification by histological analyses (Figure 7G, linear regression model p-value=0.58).

**Fig. 7.**
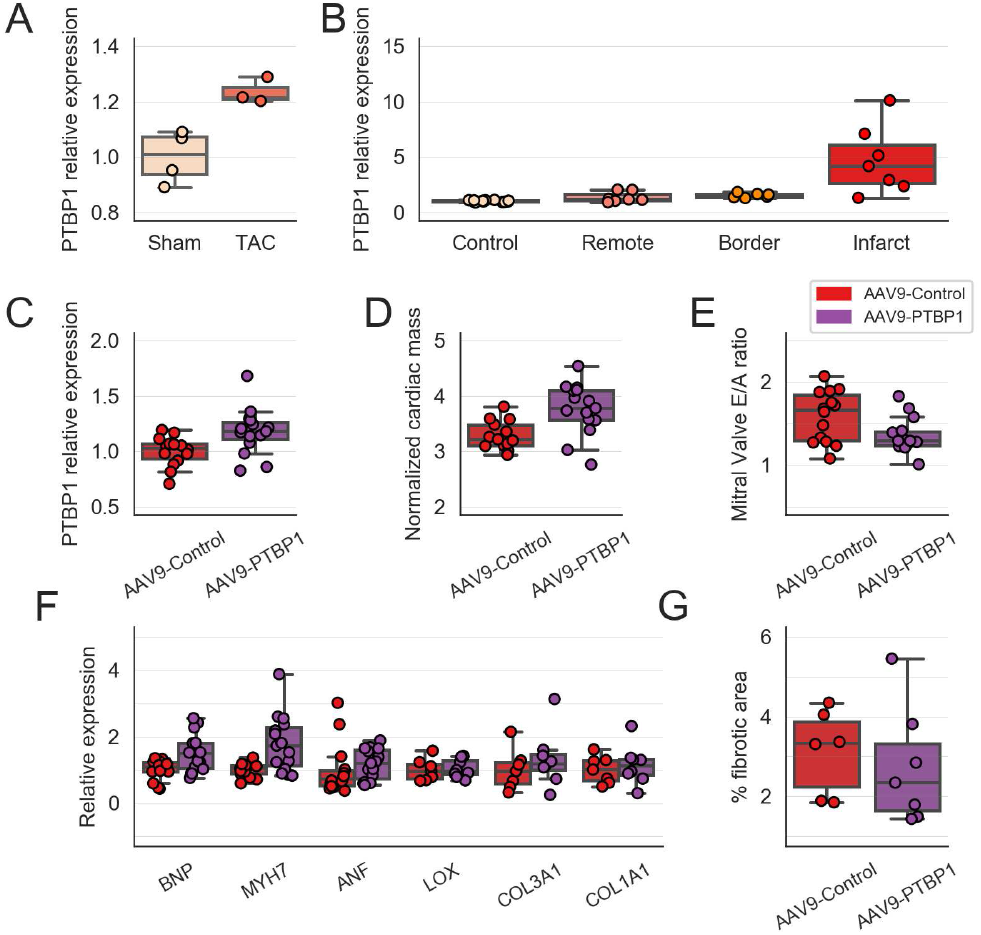
Phenotypic characterization of mice over-expressing PTBP1 using an AAV9 vector. **A** PTBP1 expression measured by qPCR in independent samples undergoing TAC (n=3) and control treatment (n=4). **B** PTBP1 expression measured by qPCR in independent samples undergoing MI (n=7), separating by infarcted, border and remote regions; and control treatment (n=13). **C** PTBP1 expression measured by qPCR in mouse hearts injected with AAV9-PTBP1 (n=15) or control virus (n=14). **D** Normalized cardiac mass derived from echocardiography analysis. **E** Ratio of E to A flow velocities through the mitral valve assessed by echocardiography. **F** Expression of markers cardiac dysfunction, hypetrophy and fibrosis measured by qPCR, normalized against GAPDH **G** Percentage of fibrotic area in histological cuts of mouse hearts (n=7 for AAV9-PTBP1; n=6 for AAV9-Control). Statistical analysis was performed using linear regression models using the treatment groups as independent variables and adjusting for batch effects when suitable. Batch effects estimated with the linear model were removed from the represented data for simpler representation

### PTBP1 over-expression partially recapitulates cardiac hypetrophy AS changes

To investigate whether PTBP1-driven cardiac hypertrophy was associated with similar AS changes to those observed in TAC at a transcriptome-wide scale, we performed RNA-seq of a reduced number of samples. The estimated *log*_2_(*FC*) in mice injected with AAV9-PTBP1 were highly correlated (Pearson *ρ* = 0.6) with those observed in TAC. Thus, PTBP1 over-expression recapitulates a great deal of the expression changes induced in pathological cardiac hypetrophy, even in absence of fibrosis (Figure 8A,B).

**Fig. 8.**
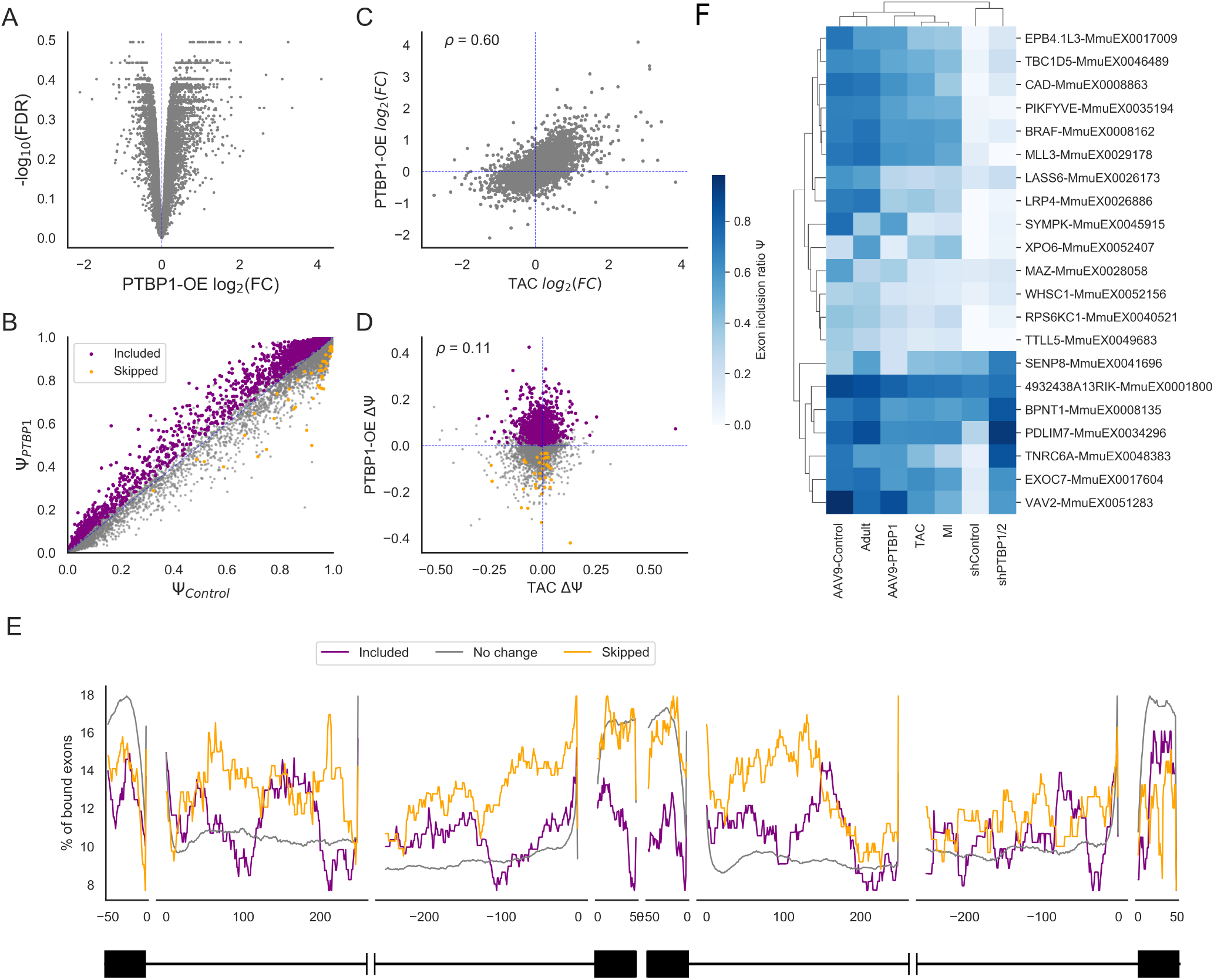
AS and GE characterization of mice over-expressing PTBP1 using an AAV9 vector. **A** Volcano plot representing adjusted p-value against *log*2 (*FC*) across all genes included in the analysis. **B** Scatter plot representing the estimated mean Ψ in hearts over-expressing PTBP1 and hearts injected with a control virus. Significant exons are highlighted in colors, depending on the direction of the change. **C, D** Scatter plot showing the relationship between the changes induced by PTBP1 over-expression (OE) and TAC for GE, assessed by *log*2 (*FC*) (C); and for AS, assessed by difference in inclusion rate (ΔΨ) (D). **E** Percentage of exons with an annotated binding site for PTBP1 by nucleotide across potential cis-regulatory regions relative to exon skipping events. Exons were classified as Included (ΔΨ > 0.1, n=329), Skipped (ΔΨ < −0.1, n=298) or unchanged (−0.1 < ΔΨ < 0.1), n=31996). X-axis coordinates represent the relative position to the closest splice site, as indicated by the scheme at the bottom. Negative coordinates represent positions upstream to the splice site. **F** Heatmap representing the estimated Ψ across conditions for high confident PTBP1-dependent cardiac hypertrophy related AS changes: exons with a ΔΨ < −0.1 after TAC, MI, AVV9-PTBP1 and ΔΨ > 0 upon PTBP1-knock down

We found a large number of exon skipping events showing significant differences between groups, mostly increased upon PTBP1 over-expression. These changes, however, showed little overlap, even at the quantitative level, to those observed in TAC (Pearson *ρ* = 0.1), similar to what we had found in PTBP1 knock-down samples (Figure S4). Even if this may be partly due to more noisy estimation of ΔΨs than that obtained for GE measures, it might well indicate that only a small part of AS changes in TAC is actually driven by PTBP1 alone (Figure 8C,D). This finding delves into the direction of a complex network of RBPs (Figure 6) underlying AS changes in MI and TAC.

To ensure that these AS changes are likely regulated by PTBP1, rather than secondary to the hypertrophy, we selected exons with an estimated ΔΨ in AAV9-PTBP1 vs control samples larger than 0.1 and lower than −0.1, as included and skipped, respectively, and plotted the distribution of CLiP-seq binding sites along relevant regulatory regions, centered at the target exon. As in TAC and MI, skipped exons showed an enrichment of binding sites in the upstream intronic flank (p-value=0.006, Table S8). We also observed a lower frequency of PTBP1 binding sites across the included exons (p-value=0.003, Table S8), suggesting that binding of PTBP1 to the target exon actually enhances its inclusion (Figure 8E). Functional enrichment analysis showed that genes with upregulated GE were mostly associated to immune response, whereas down-regulated genes were more associated to mitochondria and respiration. As before, we found different gene categories associated to AS changes: genes with included exons were associated to microtubules and muscle cell development; genes with skipped exons were weakly related to regulation of cell contraction, more directly related to cardiac function (Figure S7).

### PTBP1 regulated AS events as new candidates for understanding cardiac hypertrophy

Our results suggest that induction of PTBP1 may underlay at least some of the molecular changes associated with cardiac hypertrophy following TAC. To identify a subset of bonafide PTBP1-mediated cardiac hypertrophy AS exons, we selected a subset of exons with estimated ΔΨ < −0.1 in both TAC, MI and AAV9-PTBP1, and increased inclusion rates upon PTBP1 downregulation in neural cells (Figure 8F, Table S6), which provide interesting candidates for investigating new mechanisms underlying heart disease. Among them, we found an unannotated, but broadly conserved, 62 bp exon in intron 37 of LRP4 gene to show the largest decrease in Ψ not only in TAC and MI, but also upon PTBP1 over-expression or down-regulation (Figure S8A,B), strongly suggesting that it is PTBP1-dependent. Its inclusion increases mostly during embryonic development, and remains relatively constant during the post-natal transition. We then examined their inclusion rates across different tissues and developmental stages using VASTDB data (70) for this exon 37’ and its chicken ortholog. Indeed, its inclusion is highly specific of muscle tissues in both species, with lower inclusion rates in brown adipose tissue and mammary gland (Figure S8C,D).

## Discussion

In this study, we used an unprecedented breadth of samples and conditions to investigate the mechanisms that regulate AS changes in the heart. We found that GE and especially AS are more dynamically regulated during embryonic and postnatal heart development than after the induction of MI or TAC. Despite a partial recapitulation of developmental AS changes in heart disease, TAC and MI samples remained more similar to their adult controls than to neonatal samples, a trend also seen with GE changes. In addition, the biological processes affected by AS changes were mainly different from those altered by GE changes, regardless of the developmental or disease context, suggesting that AS and GE changes play distinct roles in the heart (23, 33, 40, 42). Interestingly, exons with increased inclusion levels during development tended to be shorter and not to affect protein domains, whereas skipped exons tended to be of similar length to unchanged exons and to encode functional domains, which would presumably be disrupted by AS, suggesting strong effects on protein function. These changes are not recapitulated in heart disease, indicating that cardiac injury triggers AS changes with a lower impact on protein function than those taking place during development.

We explored the global effect of AS changes during heart development and disease on the PPI networks. Besides AS modulated genes being central in the PPI network, as previously described in different contexts (33, 76), we found that AS changes tend to modify PPIs more often than expected by chance in both interaction datasets under study, as previously shown in cancer (9). We also analyzed the relevance of the AS-dependent interactions in the global PPI network through the edge betweenness. This property reflects how many shortest paths between nodes go through each edge, in other words, how important those interactions are for the interconnetion of different interacting modules. Our results suggest that AS changes in disease not only tend to affect more interactions than expected, but that these interactions are key in the PPI network. We did not find such association in domain mediated interactions. However, previous work suggest that most AS-mediated interactions do not affect protein domains, but short linear motifs (76). Therefore, differences between the results in the two datasets may lay on the different nature of interactions under study. These results may be limited by the small size of exons groups overlapping between the interaction datasets and the characterized exons in our study. Therefore, increasing the number of interactions or exons would help to further verify our findings. In addition to this general trend, we observed some specific AS-mediated PPI changes that may have functional impact (Table S5). Among these, we found MEF2A to have isoform specific interactions with MEOX1 and MAPK7/ERK5 and to show AS changes in both developmental transitions and MI. Interestingly, MAPK7 activates MEF2A in response to MEK5, which is alternatively spliced itself (68). Since MEOX1 and MAPK7 expression and MEF2A splicing changes are known to modulate cardiac hypertrophy (38, 44, 68), these interactions may play a role in the development of the disease. In addition, among developmentally regulated AS-mediated interactions, we found the EGFR-ERBB2-ANKS1 interaction triad to be reduced. ANKS1 regulates EGFR and ERBB2 transport to the membrane, which is necessary for its ERBB2-mediated tumorigenesis (42, 60, 71). ERBB2 deficiency has been shown to cause dilated cardiomyopathy (11), whereas its transient induction reactivates the regenerative potential of the neonatal heart in adults (17). Therefore, changes in the interaction between ANKS1 and ERBB2 or EGFR mediated by AS are expected to have major consequences for the heart.

Our results suggest that MBNL1 regulates AS in the heart mainly during development, where it has already been characterized (13, 35). MBNL1 works both as a position-dependent splicing activator and inhibitor (21). AS changes in the MBNL1 knockout mouse correlated with those observed in the ED and PD comparisons, reinforcing its main role in the regulation of AS during heart development and also validating our experimental approach. Based on these findings, we hypothesize that increased MBNL1 expression is responsible for the highly dynamic AS regulation during development, as it promotes both exon inclusion and skipping in a position-dependent manner. Although its activity may still be regulated at the protein level, the lack of enrichment suggests that its regulatory activity remains unchanged in disease, at least for the models included in this study. This may explain, at least partially, why the re-expression of neonatal patterns in disease is not complete.

PTBP1 showed the greatest contribution to explaining AS changes in heart disease and thus contribute to the partial re-expression of the neonatal AS pattern in heart disease. Whereas PTBP1 is required for the differentiation of iPSCs and fibroblasts to cardiomyocytes *in vitro* (49), and modulates the splicing of essential genes for cardiomyocyte function (e.g. Titin, Tropomyosin 1 and 2 and Mef2) (19), its role in the heart was virtually unknown. PTBP1 expression decreased during embryonic development and postnatally and was upregulated after MI or TAC. Expression changes correlated inversely with changes in the PTBP1/2 double KD, indicating that PTBP1 is a direct regulator of the AS changes taking place during development and disease. We showed how over-expression of PTBP1 in healthy myocardium is sufficient to induce cardiac hypertrophy, potentially by promoting a reduced number of the splicing changes typically induced in hypertrophic conditions, including a muscle-regulated exon in LRP4 gene. LRP4 encodes a low-density lipoprotein (LDL) receptor, which is bound by Agrin and mediates MuSK activation, which is essential for the correct functioning of the neuromuscular junction (37). In the heart, Agrin has been shown to be required for cardiac regeneration in neonates and its over-expression was able induce regeneration in adults after injury (4) and has been shown to modulate cardiomyocytes contraction *in vitro* (32). Moreover, mutations in LRP4 have been associated to sindactyly and polysindactyly in several mammalian species, including mouse, cow and humans (34, 46, 69), which is usually accompanied by cardiac defects in Timothy Syndrome patients. Inclusion of this exon introduces a stop codon near the C-terminal region of the protein and thus is expected to produce a truncated version without the last exon. Thus, even if the impact of this highly conserved and cardiac-specific splicing event in protein function remains unknown, it is an interesting candidate AS gene for better understanding of PTBP1mediated cardiac hypertrophy. PTBP1 is also known to regulate Titin splicing isoforms *in vitro* (19). Although we did not find large changes across any exon in Titin upon PTBP1 over-expression, 27 of them showed estimated ΔΨ < −0.04, which, when considered together, may contribute to isoform shortening and increased passive stiffness of the cardiac muscle as it happens during development (39).

Even if we can not rule out that PTBP1 has other functions in RNA metabolism besides AS regulation, e.g. it modulates insulin mRNA stability in the cytoplasm (22), or that AS changes are secondary consequences of PTBP1-induced hypertrophy, this seems rather unlikely, since these AS changes are associated to PTBP1 binding sites (Figure 8). Although no evidence of fibrosis was detected in mice over-expressing PTBP1, suggesting a partial recapitulation of the pathological changes observed after TAC, both mice that underwent TAC and mice injected with AAV9-PTBP1 developed diastolic dysfunction. The relevance of PTBP1 expression and function is highlighted by the fact that phenotypic changes were induced by just 1.2 fold average increase in the mRNA levels. The independent mild enrichment of PTBP2 binding sites (Figure 5A), the increased correlation coefficients observed with the PTBP1/2 double KD compared with PTBP1 single KD (Figure S4), and the simultaneous upregulation with PTBP1 in both TAC and MI (Figure 5C), suggest that PTBP2 is also playing a role in AS regulation and would further contribute to hypertrophic growth if over-expressed together with PTBP1. PTBP1 has been described as a major regulator of microexons in neurons (47). The fact that exons that undergo AS changes in the heart are shorter suggests that exon size might be a key determinant of AS regulation in heart development and disease, as is the case in the brain (33).

Interestingly, not only are the processes regulated by AS in the heart, such as vesicle transport, cytoskeleton, and cell junctions, very similar to those reported to be regulated during neural development and disease, but both tissues also seem to share sequential regulation of AS by PTBP1 and MBNL1 (9, 33, 40, 42, 74). These results suggest that AS patterns are not particularly tissue-specific, at least in brain and heart, and that they share functional and regulatory patterns that differ markedly from genes regulated at the GE level throughout development and disease.

So far, most efforts to study AS regulation have focused on the identification of specific trans-regulatory elements that, alone, can explain at least part of the AS changes (26, 35, 62). However, there is increasing evidence that AS regulatory networks are more complex: different RBPs have very similar binding affinities and bind to common targets in the RNA (14, 56, 64). How these RBPs organize in complexes and how they affect AS is not very well understood. Even considering a single RBP alone, several molecules may be required to bind stably to the target RNA, as shown specifically for PTBP1 (8), such that binding specificity is achieved by cooperative binding on nearby low affinity binding sites. Not only PTBP1 molecules bind cooperatively among themselves, but they can interact with MBNL1 proteins and cooperatively bind and modulate the inclusion of Tropomyosin exon 3 (28) or with RBM20 to regulate Titin AS (19). If interactions are widespread, a simple additive model with independent effects for all RBPs may fail to identify the actual regulatory elements, since their effects will be highly dependent on the factors that simultaneously bind each target. In this regard, we have generalized this idea by using regularized regression to allow pairwise interactions between candidate RBPs and found that such interactions are relatively abundant, and have a greater contribution to the regulation of AS in heart disease than to developmental changes. This does not necessarily imply a rewiring of the AS regulatory network, but can be explained by non-coordinated changes in the expression of the regulators. In contrast, highly coordinated changes of RBPs during development resulted in a tightly regulated set of AS changes as if they were all regulated by a single RBP, in this case, MBNL1. Moreover, by taking into account these potential interactions, PTBP1 was unveiled as an important regulator whose contribution was validated in an experimental mouse model over-expressing PTBP1. Not surprisingly, PTBP1 up or downregulation (Figures S4 and 8) alone did not reproduce a large number of AS changes compared with disease, as that would require a very specific modulation of a number of RBPs, according to our previous model. Therefore, additional studies are required to validate this hypothesis experimentally, e.g. by systematic perturbation of pairs of RBPs, and to investigate whether it is specific of cardiac disease or a hallmark of cell or tissue dysfunction.

In summary, our analysis yields biological insight about AS changes in the heart and their regulation, supported by the large number of samples included. We show that changes in GE and AS control distinct biological processes in the heart regardless of the developmental stage or disease, with AS mainly regulating actin cytoskeleton and cell junction organization. AS changes in heart disease modulate PPI more often than expected by chance, and have a strong impact on the overall PPI network. Whereas AS changes during embryonic and postnatal development are mainly modulated by MBNL1, we observed a partial re-expression of the neonatal AS pattern in heart disease likely mediated by PTBP1/2. Furthermore, changes induced by PTBP1 over-expression are sufficient to induce cardiac hypertorphy, validating our findings using a purely computational approach.

## Methods

### Dataset

We collected a series of 21 RNA-Seq experiments related to the mouse heart. Table S1 summarizes the main biological and technical characteristics of the samples used: run, experiment, pair-end, sample type, condition, sample ID, sequencer and genetic background. These experiments included samples from isolated cardiomyocytes, left ventricle, both ventricles, and the full heart at different developmental stages (embryonic, neonatal, and adult). They also included border and remote-area samples from mouse models of MI and myocardial samples from the TAC model of pressure-overload-induced cardiac hypertrophy. We filtered out samples from tissues or cell types different from those listed above as well as those from KO mice. Samples used as controls in the collected KO, TAC, or MI experiments were added to the corresponding pool of adult, neonatal, or embryonic heart samples. We additionally included data generated by our lab from infarcted mice at 7 days postinfarction (MI7d), performed as previously described (18), resulting in a total of 144 samples.

### Gene expression analysis and identification of AS events from RNA-Seq data

Gene expression (GE) and Alternative splicing (AS) were quantified using vast-tools (33). First, reads mapping to the genome were removed, and unmapped reads were later mapped to a library of exon-exon and exon-intron junctions to quantify inclusion levels for different AS event types. For GE analysis, we filtered out samples with less than 1M reads and selected as expressed genes those genes with at least 1 read per kilobase per million (RPKM) in at least 5 samples. We then used limma (65) with voom normalization to find differentially expressed genes using the experiment as random effect in the model. Since only one variable is allowed, in most cases the experiment included both tissue type and batch effect at the same time. Differentially expressed genes were defined as those with an adjusted p-value < 0.01 and an absolute *log*_2_(*FoldChange*) *>* 1. For AS analysis, we aimed to estimate the probability of inclusion of a particular event (Ψ) and to find significant differences in inclusion probabilities (ΔΨ) between conditions. We used corrected inclusion and skipping reads from vast-tools results, and selected those events supporting alternative usage (defined as having at least one read mapping to the alternative event) in at least 20% of the samples. We then used a Generalized Linear Mixed Model (GLMM) in lme4 (5) with binomial likelihood and logit link function to find differentially spliced events. As random effects, we added covariates for experiment, sample type, and individual. In this way, we added biological variability to the binomial variance in the model. We used the adult stage as baseline and stored the p-value of coefficients corresponding to the different conditions under study. Multiple test correction was applied using the Benjamini-Hochberg (BH) method. We considered as differentially spliced those events with a False discovery rate (FDR) < 0.01 and an estimated absolute ΔΨ >0.1.

### Principal component analysis (PCA)

PCA was performed using the prcomp function in R and log transformed normalized counts (adding a pseudocount) for expression. For AS, sample Ψ for each event was estimated by dividing the number of reads supporting inclusion by the total number of reads mapping to the event. We then used these estimated Ψ values as the input for PCA. Genes or exons with missing entries were removed for this analysis. All PCAs were carried out without performing batch correction to check whether the batch showed an important contribution to global variability.

### Gene ontology category analysis

Gene Ontology (GO) categories were downloaded and used as group predictors in a L1-regularized logistic regression analysis in Scikit-learn (61) to select meaningful and independent categories. We then used a standard logistic regression in statsmodels (67) to find categories with an increased probability of being represented in the selected set of genes. FDR multiple test correction was performed. We then selected the top 10 categories for each comparison and calculated pairwise semantic similarity in GOSemSim (78) using biological process (BP) ontology. A heatmap with hierarchical clustering was generated using these distances and the python seaborn clustermap function.

### Analysis of effect of AS changes on PPI networks

Domain-domain interaction (DDI) were downloaded from (24), and only pairs with corresponding human-mouse one to one orthologs were used. Mouse-human orthologs were downloaded from Ensembl-Biomart and interactions were transformed to mouse reference assuming that they will be mostly conserved. Exon domain data was downloaded from VASTDB (70), and an interaction was considered to be potentially regulated by AS whenever it was mediated by a do-main encoded by an alternative exon with that domain annotated. We then tested for an enrichment of exons mapping to domains involved in protein interactions in exons that were either skipped or included in each comparison independently. This was done using a Fisher exact test.

Isoform specific interactions were downloaded from (76). Data was collected at the isoform level, whereas our analysis was done at the exon level. To combine both, we performed the analysis at the gene level, and could not identify exon inclusion or skipping with gain or loss of interaction. Therefore, we could only test whether genes that were changing at the AS level had also isoforms showing differential protein interaction patterns. Statistical analysis was now performed at the interaction level using Fisher tests.

Each dataset was used to build an undirected graph representing AS-dependent PPI networks. We then used Networkx python library to estimate edge betweenness for each interaction. It measures how many of the shortest paths between pairs of nodes in a graph go through a particular edge. Edges connecting different modules have a high betweenness, whereas edges that do not affect network connectivity show low betweenness. To test for differences we used Mann Whitney U tests agains the unchanged group, since the distribution of betweenness was far from normal.

### CLiP-seq binding sites enrichment analysis

We extracted sequences corresponding to the 250 bp closest to the alternatively spliced exon in the flanking introns and 50 bp at both ends of the alternative and flanking exons. We then downloaded data on experimentally determined binding sites for mouse RBPs from doRiNA, Starbase, CLiPdb and ENCODE (6, 14, 45, 77). For each database, we considered any binding site detected in any sample. Overlapping binding sites for the same protein were merged using bedtools (63). CLiPdb data, provided for reference mm10, were then transformed to mm9 coordinates using the liftOver utility. Since results from the ENCODE database were obtained from human cell lines, coordinates were also transformed to mm9 coordinates using liftOver. BED files were indexed with Tabix and used to find overlaps with selected regions. We then used the one-tailed Fisher test to look for over-represented features in either included or skipped exons compared with those with no significant change. RBPs binding to specific regions showing significant enrichment (p-value < 0.01) in any of the groups of exons analyzed were subsequently used in multiple regression analysis using a GLM with logit link function and corrected for co-linearities in binding profiles.

For window-specific enrichment analysis of PTBP1 binding sites (Figure 8E), we tested the association using a Fisher Test between Included and Skipped classes, defined as having a ΔΨ > 0.1 and ΔΨ < −0.1, respectively, with the presence of PTBP1 binding sites at certain distance from each of the splice sites under analysis. We then selected the lowest p-value and the position at which the most confident enrichment was found as reported in Table S8.

### Analysis of interactions between pairs of RBPs binding sites

RBPs and regions selected for the multiple regression analysis were subsequently tested for pairwise regulatory interactions or synergistic effects. We extended the set of independent variables by adding all possible pairwise combinations among RBPs. We then used L1 regularized logistic regression using scikit-learn (61) to predict the belonging to a certain group (Included or Skipped for each comparison under study: ED, PD, TAC, MI). To select the optimal regularizing constant, we first performed 10-fold cross validation, by splitting the total number of exons in two sets, one to fit the model and one to evaluate its predictive power, over a range of regularizing constants (from 10^−3^ to 10^3^). We used the area under the Receiver Operating Characteristic curve (AUROC) curve to evaluate the predictive power for each fitting across the un-observed data in each fold. The value of the regularizing constant with highest AUROC was then selected and the model was re-fitted using all data. After model fitting, coefficient values were extracted for further analysis and classified in two types according to whether they represented single RBPs or combinations of them (also named as interactions).

### Analysis of correlation among RBPs expression levels

To calculate condition-specific correlations among pairs of RBPs we selected the samples involved in each comparison and calculated the Pearson coefficient on the log transformation of the normalized counts. Since the number and nature of samples included in each comparison is different, they were not directly comparable: correlation coefficients estimated from a reduced number of samples are expected to be noisier. We then took samples of 5000 pairs of randomly selected genes to estimate the expected mean and standard deviation of randomly selected genes, which we used to standardize correlation coefficients among RBPs. Therefore, we evaluated whether RBPs were more correlated than the expected from random genes within each group of samples. As a positive control, we calculated the z-score of the correlation for genes encoding proteins that are known to physically interact from Intact database previously used.

### Mice

Male mice housed in an air conditioned room with a 12h light/dark cycle and free access to water and chow diet were used for experiments in this study. All procedures were approved by the CNIC Ethics Committee and the Regional Government of Madrid (PA-27/13, PROEX-177/17). All animal experiments conformed to EU Directive 2010/63EU and Recommendation 2007/526/EC, enforced in Spanish law under Real Decreto 53/2013. Mice were euthanized individually using a *CO*_2_-filled chamber.

### RNA-seq analysis of PTBP1-overexpressing samples

Total RNA was extracted from the left ventricles of mice injected with the control or PTBP1 virus using the RNAeasy extraction kit (74104, Qiagen). Libraries were prepared using polyA-positive selection, and RNAs were sequenced using an Illumina-HiSeq 2500 apparatus. About 50 million paired end 100 nucleotide long reads per sample were sequenced. Generated fastq files were analyzed using vast-tools (33). Raw gene counts were analyzed with limma-voom (65) and estimated expression across samples was used for PCA. As one of the PTBP1-AAV9 samples clustered together with controls and showed a very similar PTBP1 expression, we assumed that the over-expression was not achieved in this particular mouse and was discarded from further analyses. Differential gene expression was performed using limma (65). Differential splicing analysis was limited to exon skipping events, and was performed in this case using vast-tools for pairwise comparison (33). Functional enrichment analysis was performed using GO categories downloaded from Enrichr (52) and a one-tail Fisher test. BED files containing PTBP1 binding sites were obtained from (48) and crossed with exon coordinates using Tabix. Processed data and code to reproduce figures is available at https://bitbucket.org/cmartiga/ptbp1

### Myocardial infarction and TAC surgeries

Surgeries were carried out under general anesthesia with 3-3.5% sevoflurane in 100% oxygen and providing mechanical ventilation, with a 200 μl stroke volume and a rate of 200 strokes per minute. In myocardial infarction surgery, the heart was exposed by thoracotomy, and a 8/0 silk suture thread was passed under the left anterior coronary artery and knotted to occlude the artery 1 mm distal to the left atrial appendage. In transaortic constriction, mice underwent TAC (n=3) or Sham (n=4). Thereafter, the chest wall was closed, and the skin was sutured with 6/0 nylon suture thread. Animals received analgesic treatment (0.1 mg/kg buprenorphine subcutaneously) for 3 to 5 days starting right after the surgery. Two or three days after surgery, transthoracic echocardiography was carried out to verify the successful of the procedure. Mice were euthanized 7 and 28 days after myocardial infarction surgery and 21 days after TAC surgery.

### RNA extraction and qRT-PCR

Total RNA was extracted from the LV wall using TRIzol (15596026, Thermo Fisher Scientific). First-strand cDNA was synthesized using 100 ng of total RNA and a High Capacity cDNA Reverse Transcription Kit (4368814, Thermo Fisher Scientific). Quantitative PCR (qRT-PCR) was carried out in an Applied Biosystems real-time PCR thermocycler using SYBR Green (4367659, Thermo Fisher Scientific). Primers used for qRT-PCR are listed in Table S7. Results were analyzed with LinReg PCR software (66).

### Adeno-Associated virus type 9 (AAV9) production and injection

PTBP1 and control Adeno-associated viruses were produced by the CNIC Viral Vectors Unit in HEK293 cells using serotype 9 capsid proteins. Mice were anesthetized with 3% sevoflurane in 100% oxygen. A total of 50 μL of saline (0.9% NaCl), containing 10^11^ VP/mL, were injected per mouse through the femoral vein. Two batches of experiments were performed independently: 7 control and treated mice in the first round, and 7 control and 8 treated mice in the second experiment. Mice were euthanized 28 days after injection and heart samples for RNA extraction and histological analysis were collected.

### Histological analysis

Samples were fixed in 4% paraformaldehyde in PBS during 48h, washed in PBS, dehydrated, and included in paraffin. Five-micron thick sections were rehydrated and stained using Masson’s trichrome stain. Images of each total heart section were taken under the microscope. For each sample, we analyzed the amount of fribrotic tissue by using ImageJ.

### Echocardiography

Cardiac function and chamber dimensions were analyzed by transthoracic two-dimensional (2D), M-mode (MM), and pulse wave Doppler (PW) echocardiography. Scans were performed by blinded operators using a 40 MHz linear transducer (Vevo 2100, VisualSonics). Mice were placed on a heating pad under light anesthesia with isoflurane administered with 100% oxygen. The isoflurane concentration was adjusted to maintain a heart rate of 500±50 bpm. Images were analysed by blinded operators using the Vevo 2100 analysis software (VisualSonics). Cardiac mass was normalized with the body weight. The mitral inflow pattern was assessed using PW to evaluate diastolic function from a four-chamber apical view. Diastolic function was determined by measuring the early and late diastolic peak wave ratio (E/A wave ratio).

### Accession numbers

Data generated in this study can be accessed in GEO repository (GSE119857 and GSE150831 for the the MI and PTBP1 over-expression experiments, respectively). Additional samples and datasets included in this study are listed in Table S1 and accessible in GEO repository: PRJNA141559 (43), PRJNA217955 (26), PRJNA226752 (31), PRJNA227375 (59), PRJNA239183 (20), PRJNA239462 (15), PRJNA252674 (51), PRJNA270875 (27), PRJNA270897 (57), PRJNA277489 (50), PRJNA286287 (55), PRJNA297895, PRJNA302806, PRJNA305657 (12), PRJNA306654 (10), PRJNA317318 (29), PRJNA339478 (72), PRJNA353028 (53),PRJNA379048 (16).

## Aknowledgements

We would like to thank CNIC Genomics and Bioinformatics Units for technical support and scientific discussion. We thank particularly Fernando Martínez for his help solving numerous technical problems arisen during the development of this study. Finally, we would like to thank Simon Bartlett for careful writing review of the manuscript.

## Supplementary figures

**Fig. S1.**
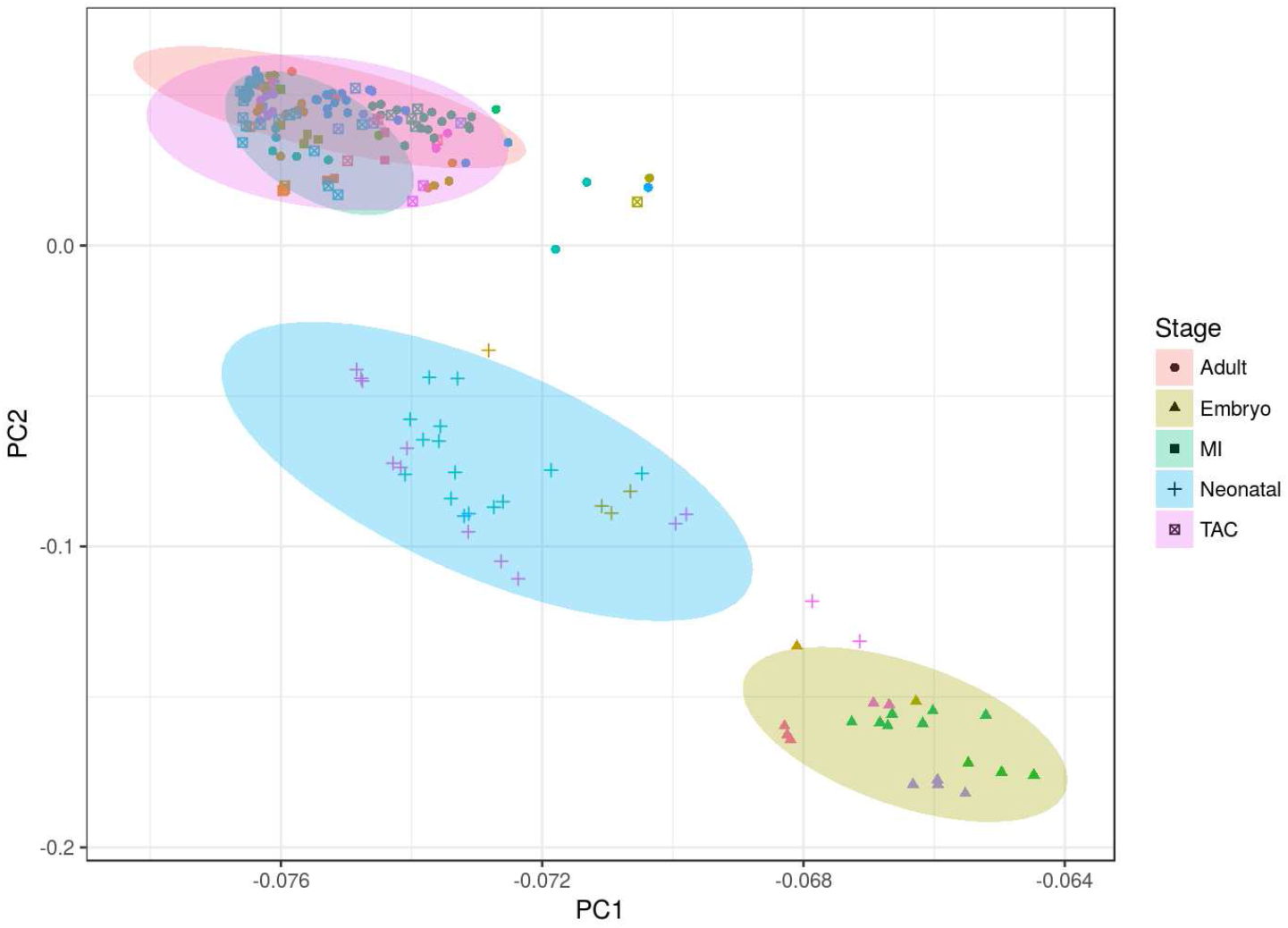
Principal Component Analysis using gene expression data. Normalized counts were transformed to z-scores, and genes with missing data were removed. Symbol shapes represent different conditions and symbol colors represent different experiments or datasets.

**Fig. S2.**
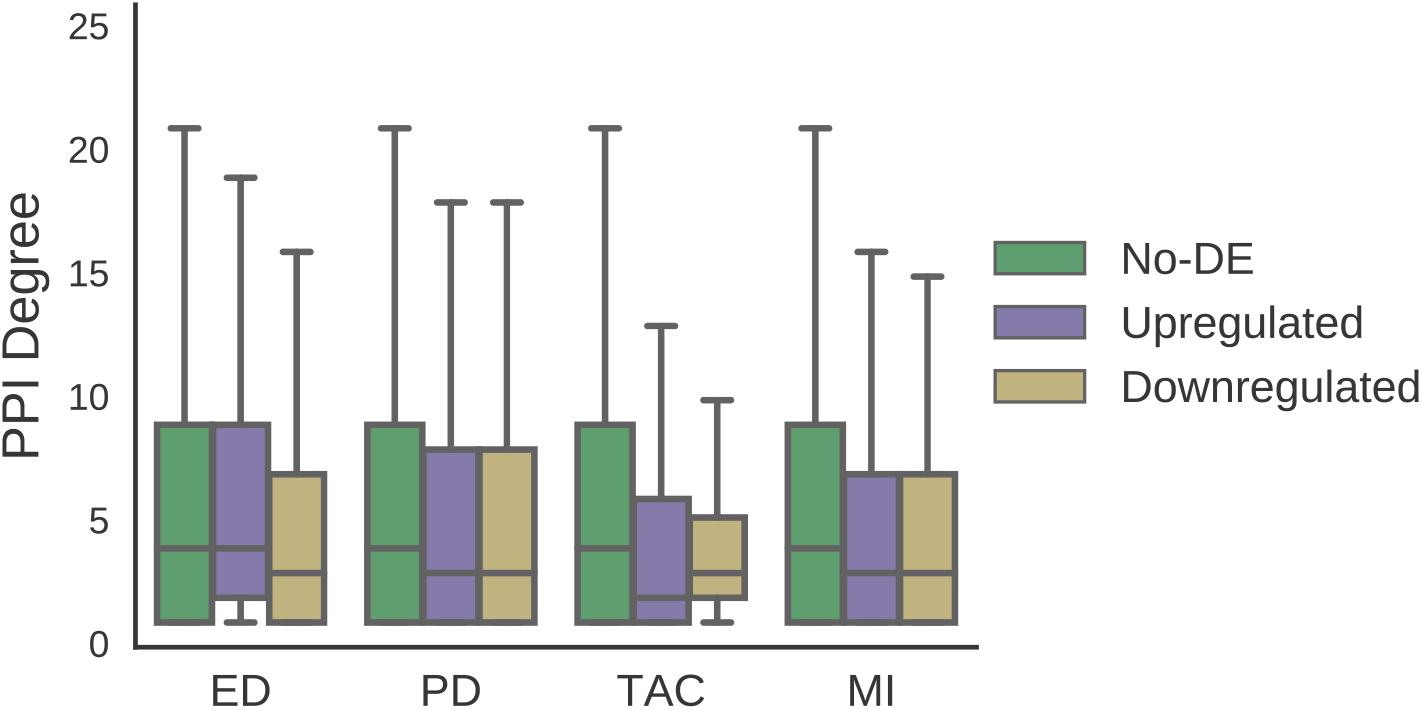
Differentially expressed genes show no changes in protein connectivity. The graph shows the degree or number of connexions in the Intact protein-protein interaction network according to whether genes were upregulated, downregulated, or remained unchanged in the comparisons studied.

**Fig. S3.**
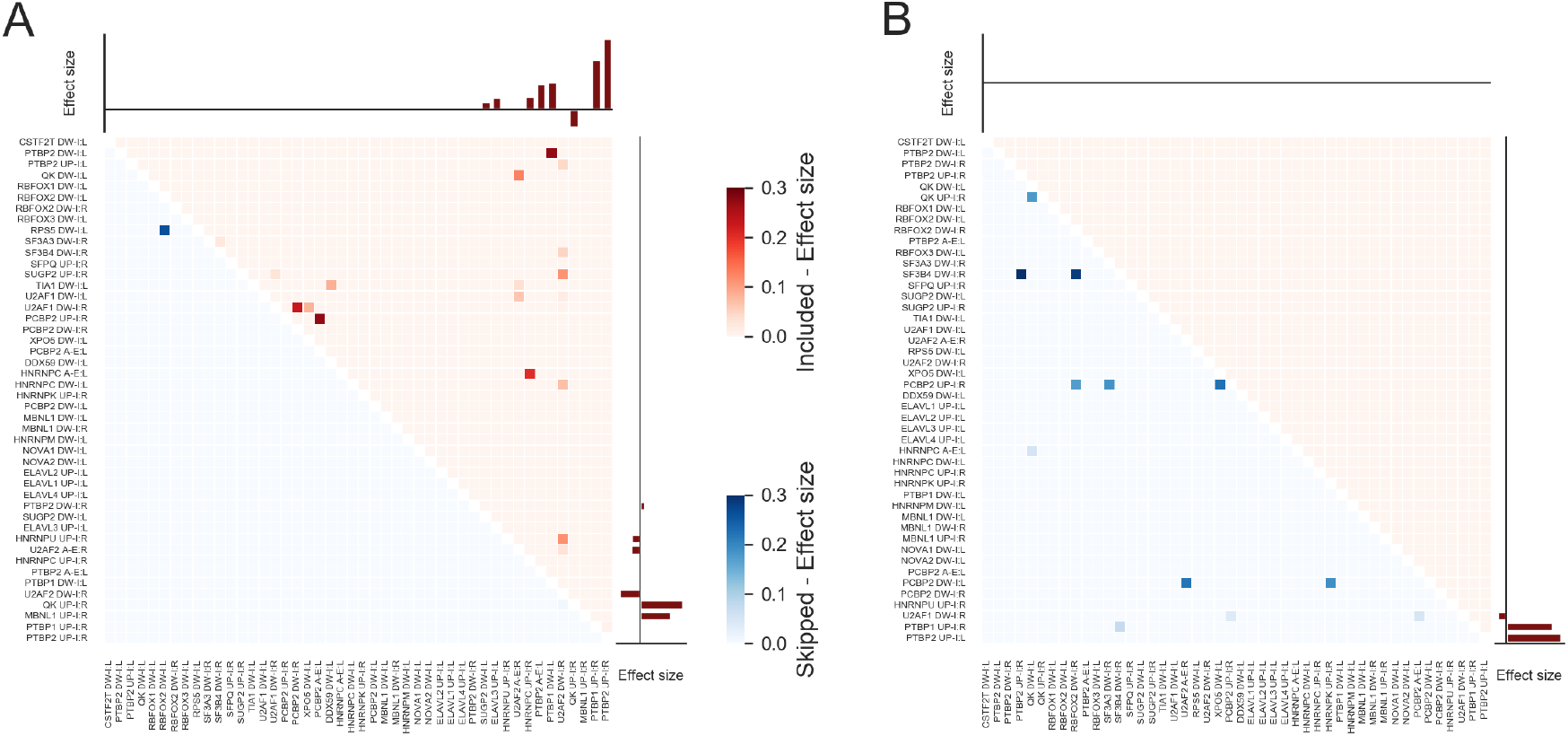
Analysis of AS regulatory complexity using regularized logistic regression with all pairwise combinations of RBPs binding sites. **A, B** Heatmap representing the estimate of the coefficients for each combination of RBPs for exons that are either included (Red) or skipped (Blue) for PD (A) and TAC (B). Barplots represent the estimation of the coefficient for single RBPs.

**Fig. S4.**
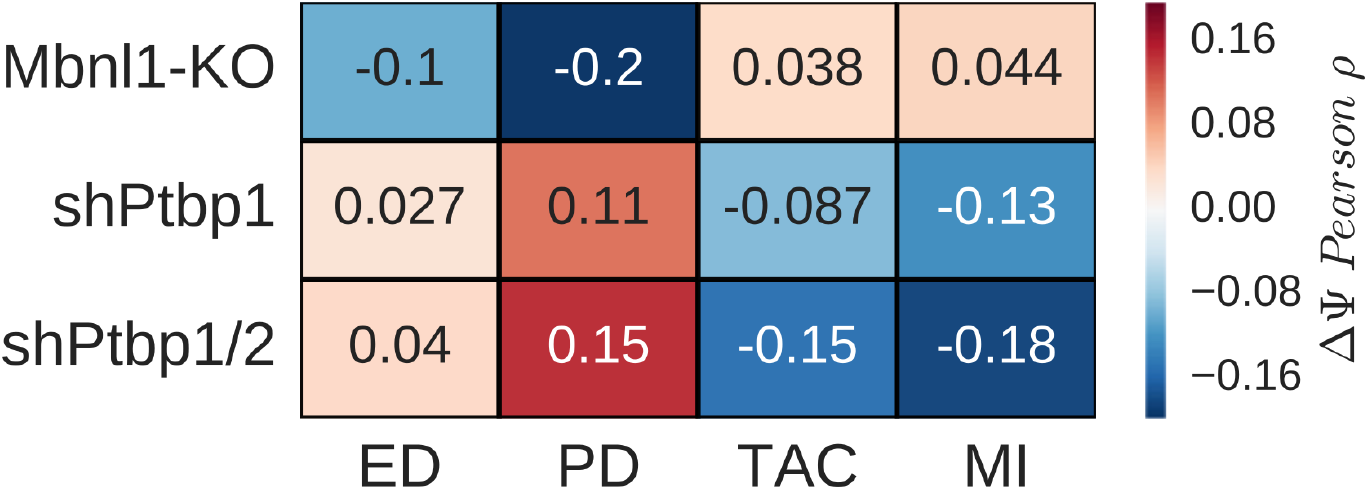
Comparison of AS changes in the heart with MBNL1 and PTBP1/2 LOF experiments. Heatmap representing Pearson correlation coefficients between changes in exon Ψ for MBNL1-KO, PTBP1-KD, PTBP1/2-KD and all comparison analyzed in the heart (ED, PD, TAC, MI).

**Fig. S5.**
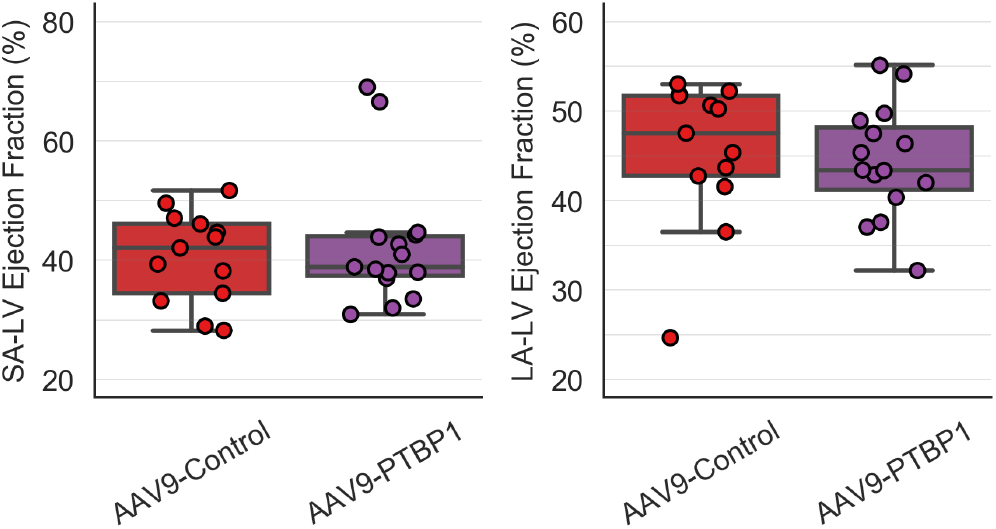
Left ventricle (LV) ejection fraction assessed by echocardiography analysis (Short Axis (SA) on the left, Long Axis (LA) on the right) in mice over-expression PTBP1 and control mice

**Fig. S6.**
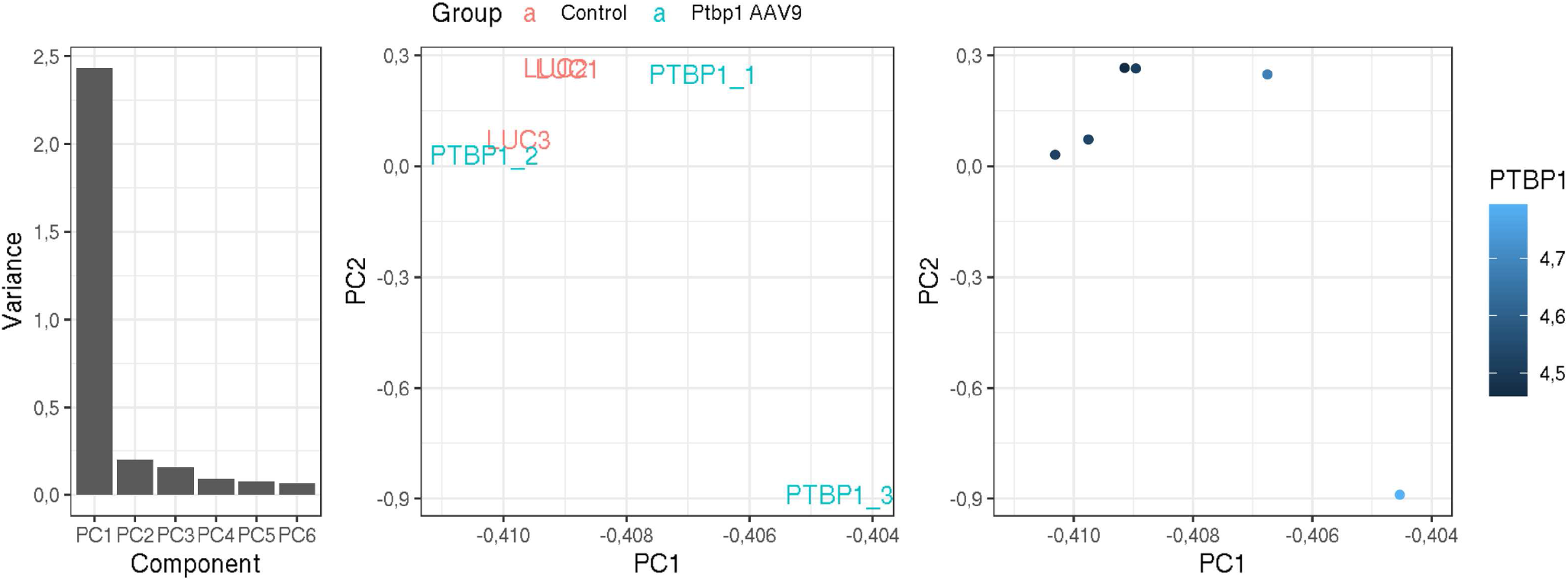
Transcriptomic characterization of mice over-expressing PTBP1 using AAV9 vector at the GE level by PCA. **A** Variance explained by each principal component. **B,C** PCA representation of samples according to the treatment group (B) and PTBP1 expression as measured in the RNA-seq (C)

**Fig. S7.**
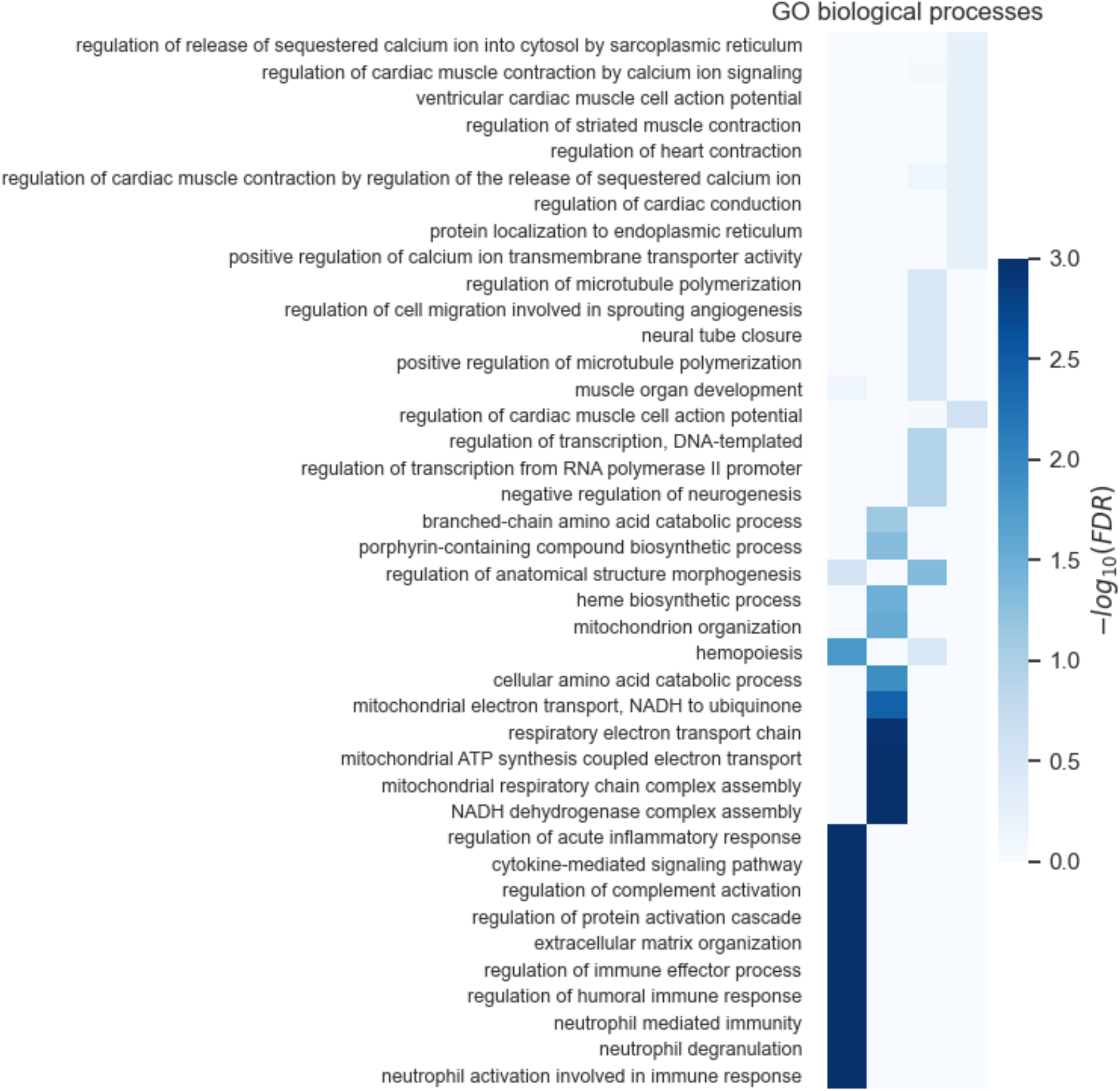
Functional characterization of transcriptomic changes induced by PTBP1 over-expression in the heart: heatmap showing the significance of the enrichment in GO functional categories, measured as −*log*(*FDR*), across different sets of genes: Up and down-regulated genes; and those showing increased or decreased exon inclusion rates (CE Included and CE Skipped, respectively)

**Fig. S8.**
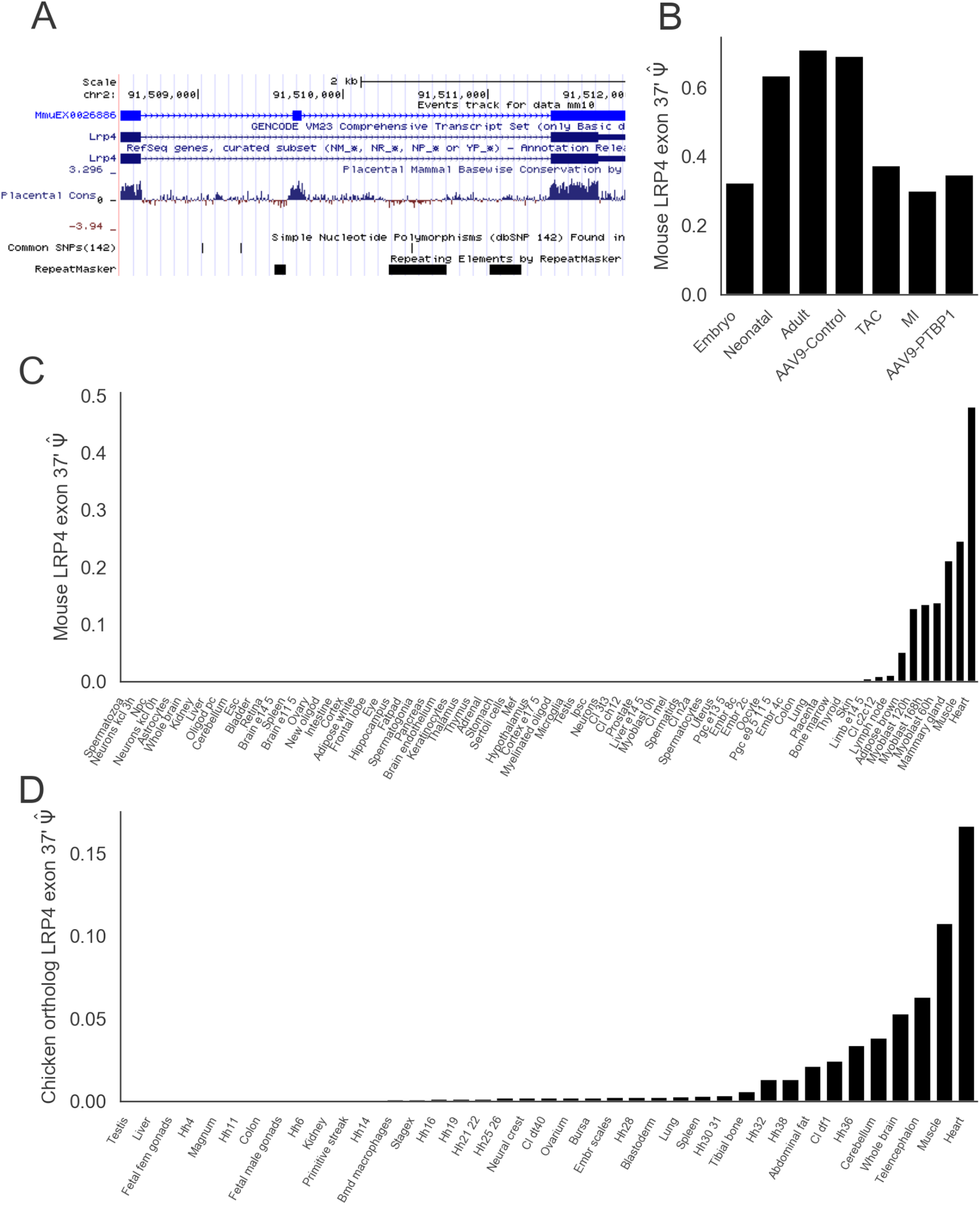
Alternative splicing of LRP4 in the heart and across tissues. **A** Genome browser snapshot of alternatively spliced exon, showing GENECODE and RefSeq annotation and mammalian conservation track. **B** Estimated exon inclusion rates Ψ across different conditions in mouse hearts. **C,D** Estimated Ψ from VASTDB (70) database for the mouse exon (C) ant its human ortholog (D) showing conserved prevalent inclusion in the heart

